# Multireplicon plasmids show broad host distributions, high transfer potential, and are enriched in antimicrobial resistance genes

**DOI:** 10.64898/2026.05.05.722934

**Authors:** Ignacio de Quinto, Paula Ramiro-Martínez, Rafael da Silva Rosa, Cristina Herencias, Val F. Lanza, Jerónimo Rodríguez-Beltrán

## Abstract

Plasmids are DNA molecules that replicate independently of the bacterial chromosome and are typically associated with the spread of antimicrobial resistance (AMR) and virulence determinants, among other relevant traits. Complex genetic rearrangements (including plasmid fusion) can result in plasmids carrying two or more replication systems, known as multireplicon plasmids. Despite decades of study, we are still far from understanding how multireplicon plasmids arise, persist, and shape the evolution of AMR.

Here, we analyzed 24,000 non-redundant plasmids across bacterial genera and found that more than 30% encoded multiple replicons. Compared with single-replicon plasmids, multireplicon plasmids were larger, enriched in antimicrobial, metal, and biocide resistance genes, virulence factors, and transfer-associated features, and observed across broader host taxonomic distributions. Some replicon pairs frequently co-occurred on multireplicon plasmids, whereas others were typically maintained on separate plasmids even when present within the same cell. Replicon pairs also showed non-random spatial organization, being preferentially located either in close proximity or at distant positions along the plasmid backbone. Their phylogenetic congruence varied substantially, suggesting that some replicon associations are evolutionarily transient, whereas others may be persistently co-inherited. Finally, we highlight the contribution of insertion sequences to multireplicon plasmid formation. Together, our work sheds light on the abundance, gene content, organization, and evolutionary dynamics of multireplicon plasmids and underscores their relevance to bacterial evolution and human health.

## Introduction

The continuous evolution of antimicrobial resistance (AMR) is outpacing human efforts to control bacterial infections^1^. Arguably, the main drivers of the development and spread of AMR are mobile genetic elements that mediate the horizontal transfer of AMR across phylogenetic boundaries^2,3^. In particular, plasmids—DNA molecules that replicate independently of the bacterial chromosome—stand out because they readily transfer AMR, virulence factors, and metabolic pathways, enabling rapid adaptation and underscoring their role as crucial agents in microbial evolution^4,5^.

Through a variety of mechanisms, including plasmid fusion, individual plasmids can acquire additional, often functional, replication systems^6–13^. These molecules, known as fusion, cointegrated, or multireplicon plasmids, are central to the alarming spread of AMR: successive fusion and fission events have transformed simpler plasmids from the pre-antibiotic era into multireplicon plasmids that pose a significant threat to resistance dissemination and, consequently, to human health^14^. The clinical relevance of multireplicon plasmids is highlighted by their remarkable ability to harbor and disseminate AMR genes, including those targeting last-resort antibiotics^15–20^. For instance, across diverse clinical settings and Enterobacterales species, multireplicon plasmids represent a substantial fraction (40-80%) of carbapenemase-associated plasmids^17,19^.

Multireplicon plasmids can emerge through at least two processes: homologous recombination between plasmids^21^, and insertion sequence (IS)-mediated plasmid fusion^22–24^. Homologous recombination can occur between repeated sequences shared by the plasmids undergoing fusion, including backbone genes, mobile elements (such as ISs), or other homologous regions^25^. In contrast, IS-mediated cointegration specifically involves transposase-catalyzed insertion of an IS-containing plasmid into a target site^26^. These pathways have been described in specific model systems, yet their relative contribution to multireplicon formation under realistic conditions remain essentially unknown.

Although multireplicon plasmids have been known for decades^27^, their origins, persistence, and role in shaping resistance evolution remain poorly understood. To address this gap, we comprehensively analyzed ∼24,000 plasmids to characterize their genetic repertoires, host distributions, structural patterns, and sequence features potentially contributing to multireplicon plasmid formation.

## Results

### Multireplicon plasmids are widespread, large, and enriched in antimicrobial, metal, and biocide resistance and virulence genes

To study the abundance of multireplicon plasmids, we retrieved plasmid sequences from PLSDB^28–30^, a curated collection of more than 70,000 high-quality plasmid genomes with associated metadata. To minimize the overrepresentation of highly similar plasmids, we deduplicated the dataset based on plasmid similarity and retained 23,925 non-redundant plasmids in which we could detect at least one replicon (**Supplementary Dataset 1**; see Methods). We found that more than 30% (7,549/23,925) of plasmids encoded two or more predicted replicons. This proportion varied substantially across host genera and, to a lesser extent, across isolation sources (**Supplementary Figures 1 and 2**): for instance, in *Klebsiella*, more than 50% of plasmids encoded two or more replicons, and nearly 25% carried three or more. In contrast, fewer than 25% of *Enterococcus* plasmids were multireplicon, and only ∼5% encoded three or more replicons (**Figure 1a**). While these estimates are subject to sequencing and annotation biases, they suggest that multireplicon plasmids are common and unevenly distributed across bacterial genera.

**Figure 1.**
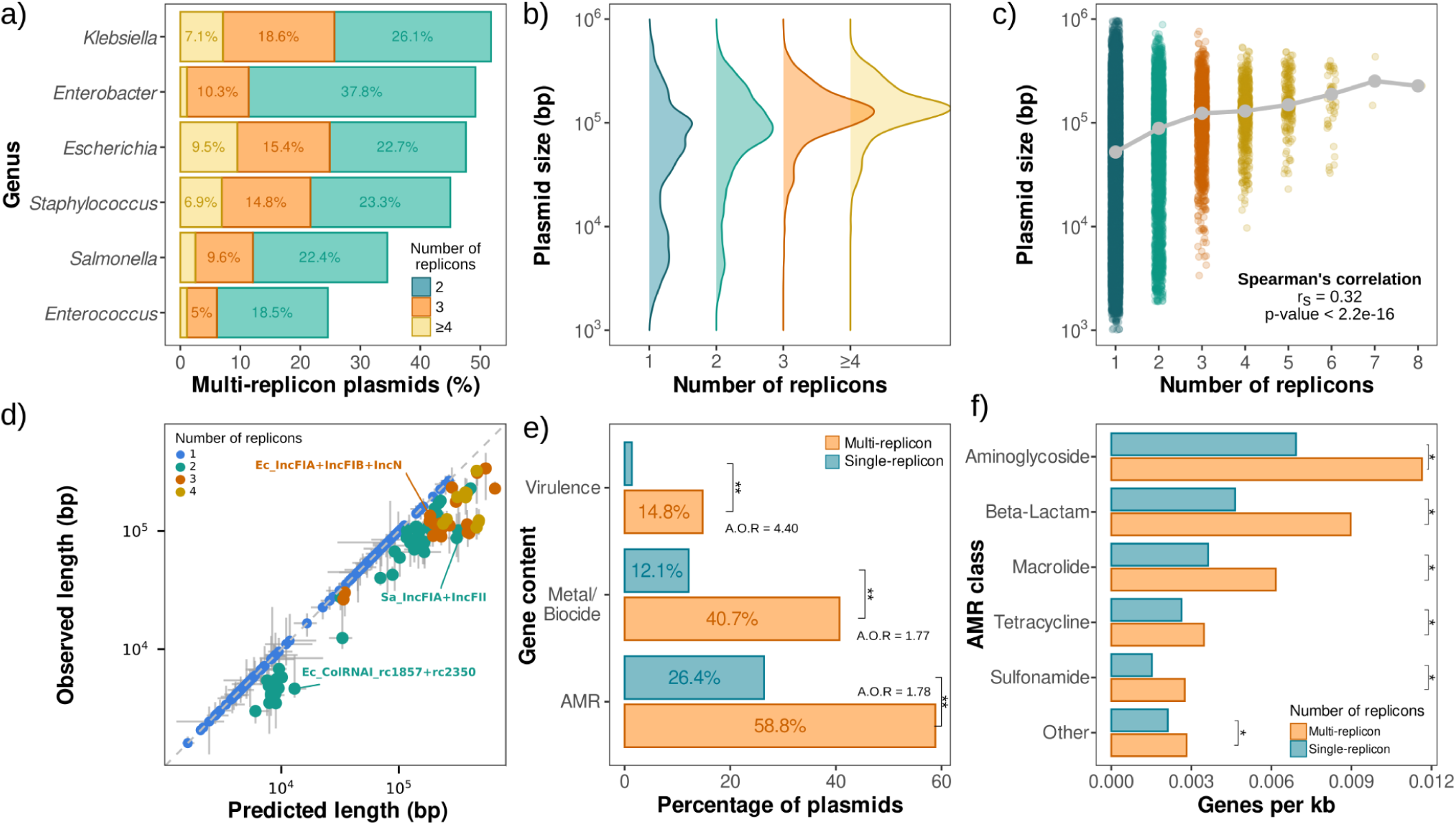
Abundance, length, and genetic content of multireplicon plasmids across genera. **a)** Percentage of plasmids classified as multireplicon (two or more replicons) in six representative genera (see legend). See **Supplementary Figure 1** for additional genera. **b)** Distribution of plasmid lengths (in bp, log scale) sorted by replicon number. **c)** Plasmid length as a function of replicon number (each point is one plasmid; y-axis on a log scale); grey points and the line represent the median plasmid length per category. **d)** Observed versus predicted plasmid length. Colors indicate the number of replicons (see legend) per plasmid. Predicted length is based on additive median lengths from single replicons, represented in blue. Points below the diagonal indicate plasmids smaller than predicted. To facilitate interpretation, some replicons are labeled. Horizontal and vertical lines show 95% confidence intervals of predicted and observed plasmid lengths respectively **e)** Percentage of plasmids encoding AMR, metal/biocide resistance, and virulence genes for single and multi-replicon plasmids. Adjusted odds ratios (A.O.R) from binomial generalized linear models controlling for plasmid length, host genus, mobility category and isolation source are shown above the bars; asterisks indicate statistical significance (Wald test p<0.001). **f)** Density of antimicrobial resistance genes for each class (genes per kb), comparing single and multi-replicon plasmids. Asterisks indicate statistical significance (Mann–Whitney U test, p < 0.001 in all cases).

Multireplicon plasmids may arise through multiple mechanisms, including the acquisition of additional replication modules and plasmid fusion. Fusion can generate larger molecules that may subsequently undergo genomic rearrangements and size reduction^14,31^. Consistent with this, the length distribution of multireplicon plasmids was skewed toward larger plasmids, with the canonical bimodal plasmid length distribution^32^ becoming unimodal for plasmids carrying more than three replicons (**Figure 1b, Supplementary Figure 3**). Across the most represented bacterial genera in the dataset, plasmid length was positively correlated with the number of replicons, albeit only modestly (Spearman’s ρ = 0.32, p < 0.001; **Figure 1c, Supplementary Figure 4**). Moreover, multireplicon plasmids were typically smaller than the summed median lengths of single-replicon plasmids carrying their constituent replicons (**Figure 1d**). Together, these observations are compatible with plasmid fusion followed by reductive evolution, although alternative mechanisms cannot be excluded^14,31^.

We then examined gene content, focusing on AMR, metal and biocide resistance, and virulence genes, as these clinically relevant functional categories are frequently plasmid-borne^4,33,34^. For each category, we estimated the proportion of plasmids carrying at least one gene in that class. In agreement with previous studies^14,35^, AMR genes were detected in nearly 60% of multireplicon plasmids but in only 26% of single-replicon plasmids. Similarly, metal/biocide resistance genes were found in 40% of multireplicon plasmids and only 12% of single-replicon plasmids, and virulence genes in 15% and 1%, respectively (**Figure 1e**). All differences were statistically significant after controlling for plasmid length, predicted mobility, host genus, and isolation source using binomial generalized linear models (Wald test; adjusted odds ratio = 1.78, 1.77, and 4.40, for AMR, metal/biocide and virulence, respectively; p < 0.001 in all cases; **Supplementary Table 1**). Importantly, this enrichment was not simply explained by their larger size, as multireplicon plasmids also carried higher densities of AMR, metal/biocide resistance, and virulence genes than single-replicon plasmids (**Figure 1f, Supplementary Figure 5**; Mann–Whitney U test, p < 0.001 in all cases).

### Multireplicon plasmids are enriched in mobility-associated features and are associated with broader replicon host distributions

We next compared multireplicon and single-replicon plasmids across two crucial plasmid traits: mobility and taxonomic distribution. Mobility-associated features were consistently more frequent in multireplicon plasmids, even after adjustment for plasmid length, host genus, and isolation source (**Supplementary Figure 6**). Using a four-category classification based on the detected presence of mobility components^36–38^, 47% of multireplicon plasmids were predicted to be conjugative (pCONJ), compared with only 26% of single-replicon plasmids even after adjusting for plasmid length, host genus and isolation source (adjusted OR = 1.22, 95% CI: 1.13–1.31). Conversely, presumably non-transmissible plasmids (pNT) were less frequent among multireplicon plasmids than among single-replicon plasmids (19.4% vs. 38.9%; adjusted OR = 0.50, 95% CI: 0.46–0.53; **Figure 2a, Supplementary Table 2**). Although these results were broadly supported by independent prediction tools (**Supplementary Figure 6 and 7**), we note that sequence-based predictions do not necessarily reflect functional transfer capacity.

**Figure 2.**
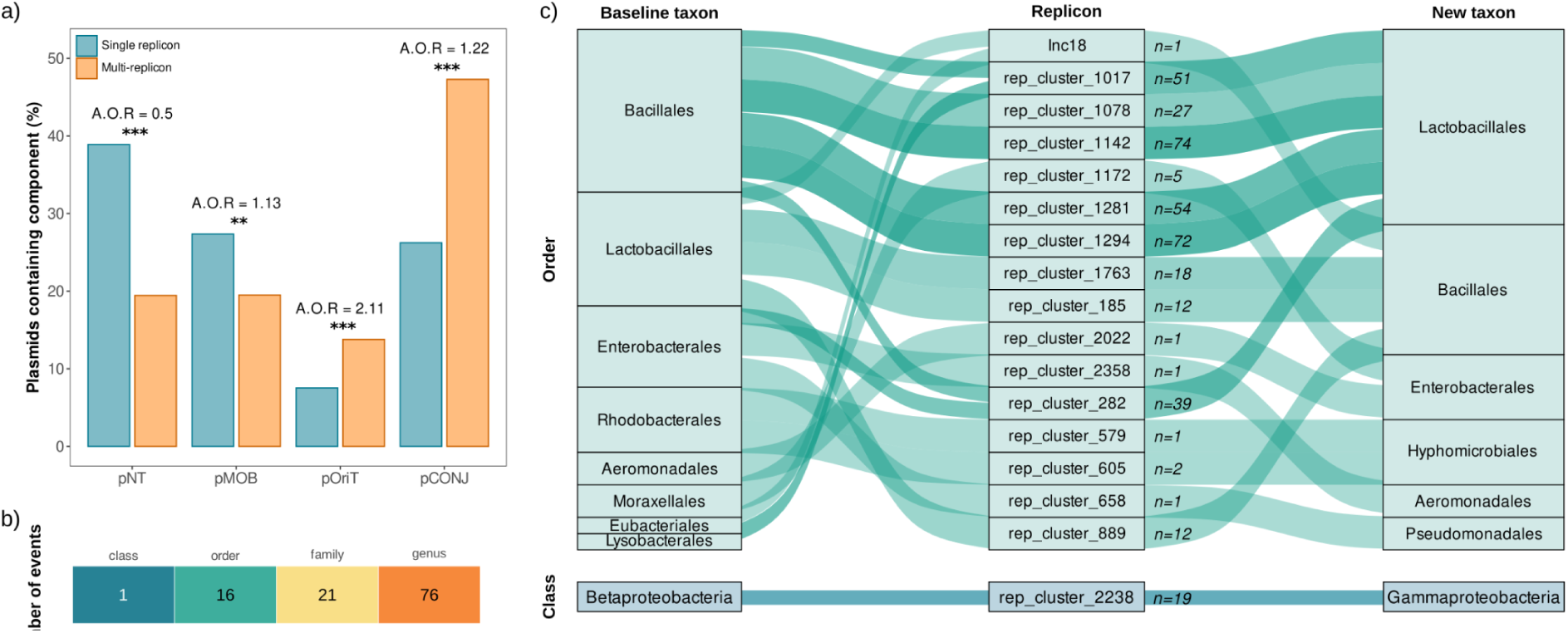
Multireplicon plasmids show higher mobility potential and broader observed host distributions. **a)** Percentage of single-replicon (n = 16,376) and multireplicon (n = 7,549) plasmids assigned to four mobility categories based on their predicted transfer machinery: conjugative (pCONJ, encoding both a relaxase and a mating-pair formation system), mobilizable (pMOB, encoding a relaxase but lacking a complete mating-pair formation system), oriT-containing (pORIT, carrying a predicted origin of transfer but lacking a relaxase and a complete mating-pair formation system), and non-transmissible (pNT, lacking identifiable transfer-associated components). Adjusted odds ratios (A.O.R) from binomial generalized linear models controlling for plasmid length, host genus, and isolation source are shown above the bars. Asterisks indicate statistical significance (Wald tests, p < 0.01) n.s. denotes non-significant differences. Notably, although the observed proportion of pMOB was lower among multireplicon plasmids, this association reversed after adjustment (AOR = 1.13). **b)** Number of replicon-taxa combinations that showed broader observed host distributions in multireplicon plasmids at different taxonomic levels. **c)** Replicons showing broader host range as multireplicon plasmids at the order and class levels. Baseline taxa in which the focal replicon was observed on single-replicon plasmids are shown on the left, focal replicons in the center, and additional taxa in which they were observed on multireplicon plasmids on the right. Labels indicate the number of observed assemblies for each event. Because one replicon may be present as a single replicon in multiple baseline taxa it can contribute multiple visualization edges in the left part of the plots, but is counted only once in panel b. Events at the family rank are shown in **Supplementary Figure 8**.

We next asked whether individual replicons were found across a broader range of bacterial hosts when associated with other replicons in multireplicon plasmids. To investigate this, we analyzed the complete, non-deduplicated PLSDB dataset of 72,556 plasmids. For each replicon, we established a baseline host distribution comprising the genera in which it occurred in single-replicon plasmids. We then identified additional genera in which the same replicon was detected in multireplicon plasmids but not in single-replicon plasmids. This analysis identified 289 cases in which a replicon was found exclusively in multireplicon plasmids in a genus beyond its observed single-replicon host distribution (**Supplementary Dataset 2**).

To independently evaluate these associations using a larger genomic dataset, we analyzed 1,863,797 bacterial assemblies from the 78 implicated genera in the AllTheBacteria database^39^. We reasoned that, if these replicons showed broader host distributions when present in multireplicon plasmids, the focal replicons should not be detected alone in the newly observed host genera, and should instead co-occur with their corresponding partner replicons. This led to the exclusion of 66 cases in which the focal replicon was detected without any other identifiable replicon in at least one assembly from the new genus, suggesting that its occurrence in that genus might not be restricted to multireplicon plasmids. For the remaining cases, we required the focal replicon to co-occur with at least one of its PLSDB partner replicons in at least one assembly from the additional genus. Although such co-occurrence does not establish that the replicons reside on the same plasmid, it provides independent evidence consistent with the replicon combinations observed in PLSDB. Under these stringent criteria, we identified 114 cases in which a replicon was detected outside its baseline single-replicon host distribution in association with the partner replicon identified in PLSDB. Of these, 76 involved additional genera, 21 extended the observed distribution to additional families, 16 to additional orders, and one to an additional class. These cases were represented in a median of 7.5 assemblies, with 26 represented by a single assembly and 88 by at least two (**Figure 2b–c, Supplementary Figure 8**).

### Non-random association of replicons in multireplicon plasmids

To study the patterns underlying multireplicon assembly, we analyzed a subset of 13,237 multi-replicon plasmids from six clinically relevant bacterial genera spanning Gram-negative Enterobacterales (*Klebsiella, Enterobacter, Escherichia,* and *Salmonella*) and Gram-positive Firmicutes (*Enterococcus* and *Staphylococcus*). We constructed a replicon association network in which nodes represent replicons and edges link replicon pairs observed on the same multireplicon plasmid (**Figure 3a**). The network comprised two largely independent clusters, corresponding to Enterobacterales and Firmicutes, with the Enterobacterales cluster being markedly more interconnected (**Supplementary Dataset 3**). Within each module, several replicons behaved as hubs, co-occurring with many partners. In Enterobacterales, IncF-like replicons (IncFIA, IncFIB, IncFIC, IncFII) were highly interconnected among themselves and with other common replicons, such as IncHI1B, IncR, and IncQ1. Notably, a small number of replicons also served as bridges between the Enterobacterales and Bacillales clusters. For instance, Firmicutes-associated Inc11, rep_cluster_1018, and rep_cluster_282 were observed in combination with Enterobacterales-associated rep_cluster_2350, IncFIC, IncP, or IncR (**Figure 3a**, **Supplementary Dataset 3**). This is consistent with our previous observation that replicons typically found in distantly related hosts can occur together in multireplicon plasmids.

**Figure 3.**
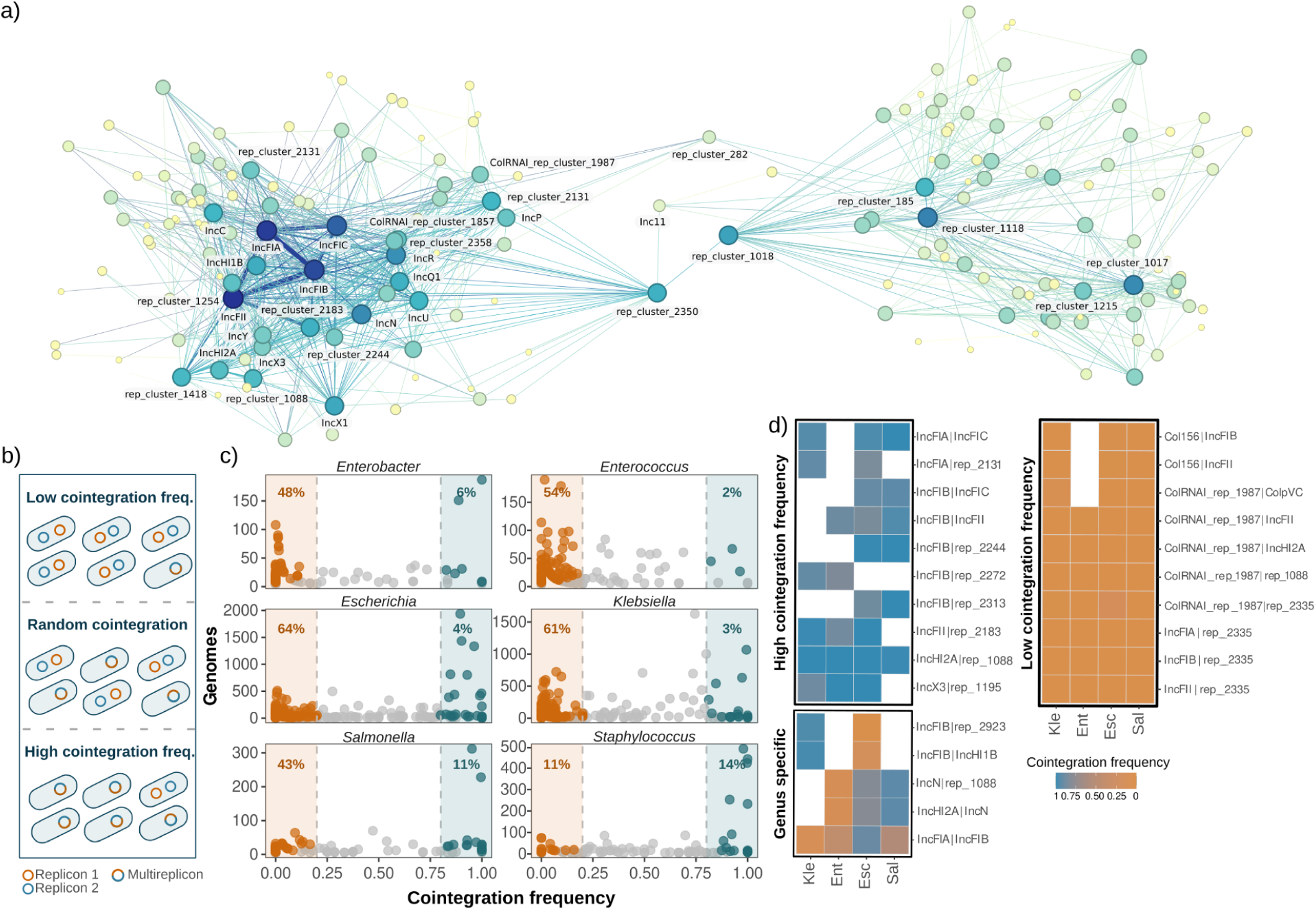
Co-occurrence patterns of multireplicon plasmids. **a)** Replicon association network. Each node represents a replicon, and edges connect pairs of replicons that co-occur within the same multireplicon plasmid. Node size is proportional to the number of co-occurring partners (degree), and edge width reflects the frequency of multireplicon plasmids for a given replicon pair. **b)** Conceptual framework for cointegration frequency. Schematic representation of low, random, and high cointegration frequencies between two replicons (blue and red circles). **c)** Genus-specific distribution of cointegration frequencies. Scatter plots show cointegration frequencies for replicon pairs within each genus. Each point represents a replicon pair. The percentages of high– and low-cointegration frequency interactions are shown in each panel. **d)** Conserved and genus-specific associations. Heatmaps display replicon pairs and their cointegration frequencies across genera (see legend).

Because prevalent replicons in the dataset may appear disproportionately connected within the association network (e.g. IncF-like replicons in *Klebsiella* and *Escherichia*), we next quantified co-occurrence at the cellular level. Specifically, for replicon pairs coexisting intracellularly in at least five distinct genomes, we calculated their *cointegration frequency*: the tendency to reside on a single multireplicon plasmid rather than two separate ones (**Figure 3b**). We identified 73 replicon pairs that were preferentially observed together on multireplicon plasmids (cointegration frequency ≥ 0.8) and rarely co-occurred as independent plasmids in the same cell (though they may be abundant individually). In contrast, 1,078 replicon pairs preferentially co-occurred as independent plasmids and were only rarely observed together on a multireplicon plasmid (cointegration frequency ≤ 0.2). The remaining 896 pairs either showed intermediate cointegration frequency (0.2 < cointegration frequency < 0.8) or could not be confidently classified due to limited statistical power (**Figure 3c, Supplementary Dataset 4**). Cointegration frequencies remained quantitatively consistent even after balancing the disproportionate contribution of overrepresented genus-replicon combinations (**Supplementary Figure 9**). Nevertheless, these results are influenced by the taxonomic and replicon composition of the dataset, limiting their generalizability to underrepresented taxa and replicons.

Although the proportion of high and low cointegration frequency replicon pairs differed by genus (**Figure 3c, Supplementary Dataset 4**), these replicon pairs generally showed consistent cointegration frequencies (**Supplementary Figure 10**). For instance, several IncF-associated pairs repeatedly showed high cointegration frequencies, whereas multiple Col-like replicons (ColRNAI, Col156, and rep_cluster_2350) showed consistently low cointegration frequency with larger plasmids, particularly those of the IncF family (**Figure 3d, Supplementary Dataset 4**). Despite this overall conservation, we occasionally observed that certain replicon pairs were frequently found on multireplicon plasmids in one genus but preferentially occurred on separate plasmids in another (**Figure 3d**). Remarkably, *Escherichia* and *Klebsiella* showed contrasting patterns in various replicon pairs, most of which included the IncFIB replicon.

Altogether, these results show that replicon associations within multireplicon plasmids are non-random and cannot be explained by within-cell co-occurrence alone. We note, however, that cointegration frequency should not be interpreted directly as replicon fusion propensity, because it may reflect both the formation and the subsequent persistence of a given replicon association.

### Conserved replicon spacing is associated with phylogenetic congruence

We next examined the distance between replicons in multireplicon plasmids. We reasoned that if plasmids merge randomly, the distances between pairs of replicons would be uniformly distributed. To evaluate this, we calculated the *relative topological distance* for each replicon pair in our dataset, defined as the shortest distance between two replicons in base pairs, normalized by the total length of the multireplicon plasmid. For circular plasmids, relative distances range from approximately 0 for adjacent replicons to 0.5 for replicons located at diametrically opposite positions on the plasmid (**Figure 4a**).

**Figure 4.**
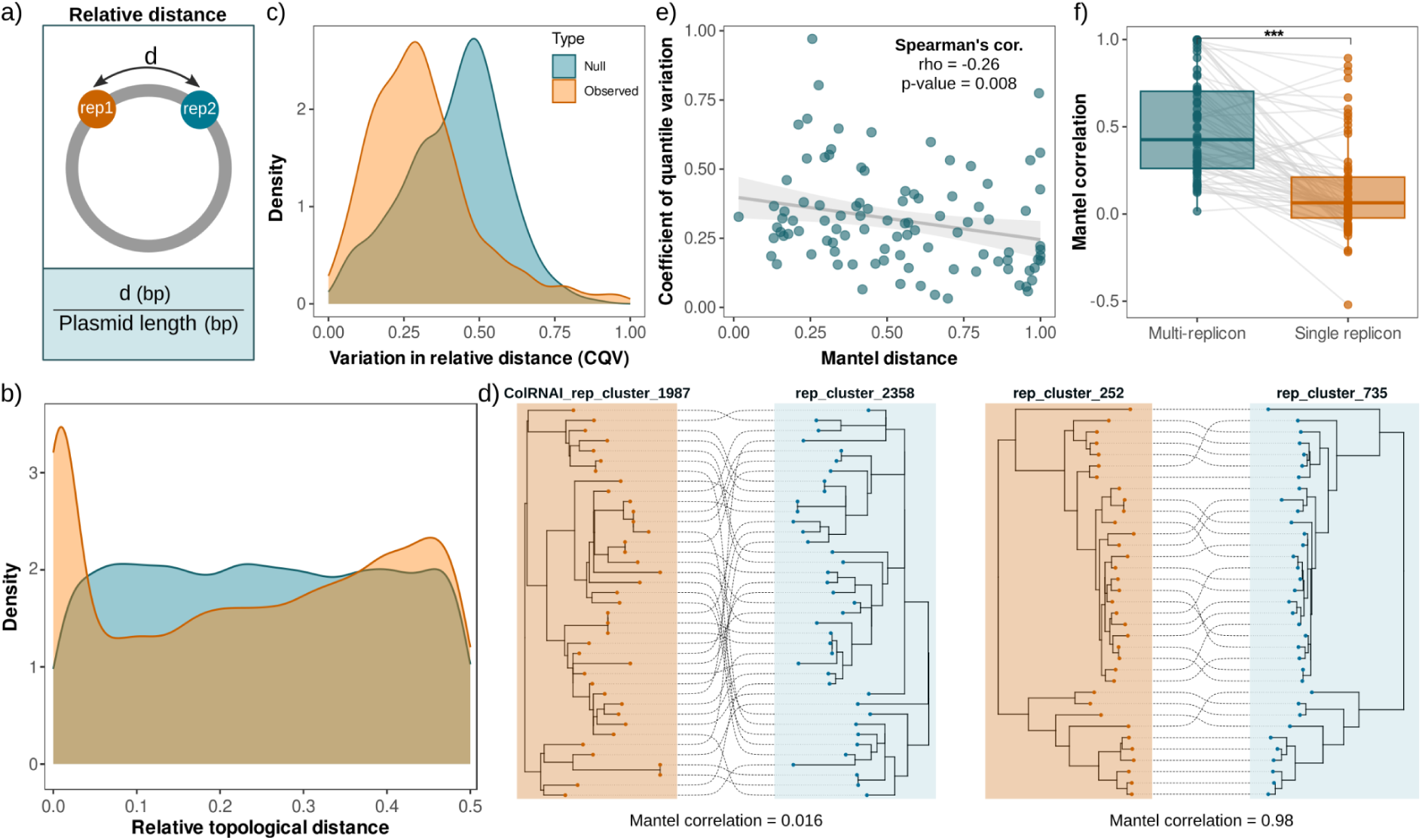
Replicon spacing and phylogenetic congruence. **a)** Relative topological distance between replicons. The minimal distance (d, in bp) separating two replicons (rep1 and rep2) is normalized by total plasmid length to obtain the relative topological distance. **b)** Density plots compare distances between replicon pairs (observed; orange) with a null model (null; blue) that assumes random positioning along the plasmid backbone. **c)** Density distribution of the quantile coefficient of variation (CQV) for relative distances between replicon pairs. Observed data (orange) show a shift toward lower variability compared to the null expectation (blue), indicating that relative topological distances are more conserved than expected by chance (Mann–Whitney U test, p < 10⁻^10^). **d)** In these representative examples, tanglegrams compare the phylogenetic trees of two replicons in multireplicon plasmids that carry both. An entangled tanglegram (Low Mantel coefficient, on the left) indicates limited phylogenetic congruence, whereas an untangled tanglegram (high Mantel coefficient, right) indicates stronger phylogenetic congruence. **e)** The scatter plot shows the association between variability in relative distances (CQV) and phylogenetic congruence (Mantel correlation). Each point is a pair of replicons. Replicon pairs with more conserved spacing exhibit stronger phylogenetic congruence. **f)** Phylogenetic congruence in single vs multireplicon plasmids. Boxplots show the Mantel correlation coefficient for replicon pairs in multireplicon plasmids vs. in two separate single-replicon plasmids. Points represent individual replicon pairs, and gray lines connect matched pairs occurring together in multireplicon plasmids or separately in single-replicon plasmids. Asterisks denote statistical significance (paired Wilcoxon signed-rank test p < 10^-10^.

We found that, compared with the random null model, the distribution of relative topological distances was markedly non-uniform (**Figure 4b**; Kolmogorov-Smirnov test p < 10⁻^13^). Relative topological distances peaked near 0 and, to a lesser extent, near 0.5, with fewer observations at intermediate values. This indicates that replicon pairs tend to cluster either closely or at opposing positions along the plasmid sequence, rather than being randomly distributed.

This non-random organization suggests that specific replicon pairs preferentially associate at defined relative distances. Consistently, we observed that the relative topological distance between most replicon pairs was relatively constant, displaying low variation (one-sided Mann–Whitney U test, p < 10⁻^10^; **Figure 4c**), suggesting that most replicon pairs conserve a relatively stable topological distance across individual plasmids.

We next asked whether conserved replicon spacing was associated with phylogenetic congruence between replicon pairs, as might be expected if they shared evolutionary history. To test this, we built a separate phylogenetic tree for each replicon and quantified the correspondence between paired phylogenies using Mantel tests. High Mantel correlation coefficients indicate strong phylogenetic congruence, whereas Mantel coefficients close to zero indicate little congruence^40,41^(**Figure 4d**). Replicon pairs with more conserved relative spacing showed greater phylogenetic congruence (Spearman’s ρ = −0.27, n = 97, p < 0.01, **Figure 4e**), regardless of whether they were located close together or far apart along the plasmid (**Supplementary Figure 11**). Notably, phylogenetic congruence was significantly lower when the same replicon pairs coexisted as separate plasmids (**Figure 4f**). Together, these results suggest that once combined into the same multireplicon plasmid, some replicon pairs can be co-inherited, thereby preserving their relative positions and showing shared evolutionary histories. Furthermore, the regions flanking replicons with high Mantel coefficients also tended to show congruent phylogenies (Spearman’s ρ = 0.269, **Supplementary Figure 12**), suggesting that shared evolutionary history may extend beyond the replicons themselves. Conversely, low congruence between replicon pairs may reflect recurrent independent fusion events or the erosion of a shared evolutionary history through recombination and replicon turnover.

### Breakpoint sequences are consistent with homologous recombination and IS-associated fusion signatures

Several mechanisms may contribute to multireplicon formation. We focused on two well-described routes of plasmid fusion, homologous recombination and insertion sequence (IS)-mediated cointegration, because both leave direct sequence duplications at the junctions between the parental plasmids, providing a detectable molecular signature of fusion^21,23,24,42^.

To identify candidate fusion events generated through these routes, we searched the database for cases in which a smaller, independent plasmid (containing fewer replicons) was embedded within a larger multireplicon backbone by aligning all multireplicon plasmids against all possible smaller plasmids. We retained only continuous hits where over 90% of the smaller plasmid sequence (at >90% nucleotide identity) was embedded in a larger multireplicon plasmid. Using this approach, we successfully identified 1,617 multireplicon plasmids in which at least one smaller plasmid was contained, enabling us to map the precise junction breakpoints.

We then extracted 500 bp flanking regions on either side of the identified junction breakpoints and aligned them against the corresponding opposite junction within the same multireplicon plasmid to search for molecular signatures of plasmid fusion (**Supplementary Figure 13**). We found that nearly 70% of the plasmids exhibited no detectable (29%) or very short homology stretches (42%), below the minimum length required for efficient RecA-mediated recombination (<20 bp at >95% identity; **Figure 5a**)^43^. This suggests that these structures either arose through alternative routes, such as RecA-independent mechanisms, or that the signatures of homologous recombination or IS-mediated cointegration have been eroded beyond detection.

**Figure 5.**
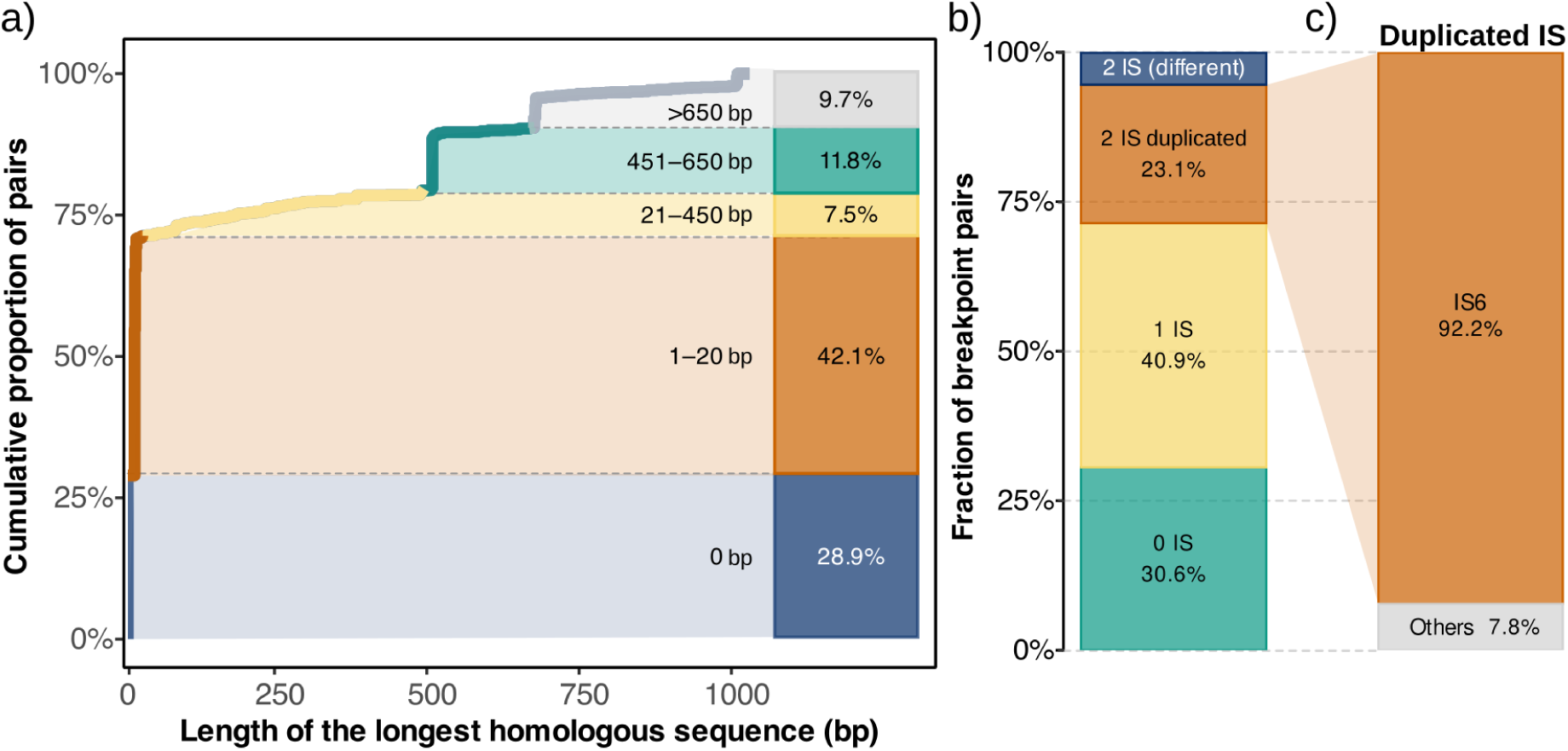
Sequence homology and insertion sequence patterns at inferred plasmid junctions. **a)** Cumulative distribution of the longest homologous sequence detected between breakpoint regions, grouped by length. The stacked bar plot summarizes the proportion of window pairs by homology length categories. For each category, the base-pair (bp) range and relative proportions are shown. **b)** Fraction of breakpoint pairs classified according to insertion sequence (IS) content, including no IS, one IS, two IS (duplicated or different), highlighting the diversity of structural contexts associated with fusion. **c)** Proportion of IS families in cases where the same IS was found in both breakpoints.

In contrast, 29% of multireplicon plasmids (469/1,617) showed larger homology regions (>20 bp at >95% identity) within the analyzed windows, consistent with fusion involving shared homologous sequences. To assess whether such homology was common elsewhere in these plasmids, we performed pairwise alignments of length-matched random windows from the same plasmids and found little to no detectable homology (**Supplementary Figure 14**). Notably, the cumulative distribution of the homologous tracts exhibited sharp increases at specific lengths, particularly around 500 bp and 650 bp (**Figure 5a**), suggesting the recurrent involvement of sequences of similar lengths.

To explore the nature of these sequences, we grouped the identified homologous regions into clusters based on DNA similarity and analyzed their genetic content. Although most sequences (78%) were unique or found in fewer than 15 plasmids, a single cluster accounted for 22% of all sequences (**Supplementary Figure 15**), suggesting that a conserved sequence may contribute to a substantial fraction of the putative fusion events. Sequence similarity searches revealed that this frequent cluster partially matched IS6 family members.

Prompted by this observation, we next searched IS elements present at both junctions using expanded window sizes of 2kb to account for the known variability in IS lengths^24^(**Supplementary Figure 16**). We also searched for IS elements in randomly selected, equivalent-sized windows across plasmid sequences to control for basal IS prevalence (**Supplementary Figure 17**).

Using this approach, 23% of candidate fusion events exhibited matching IS annotations at both junctions (**Figure 5b**), compared with only 3.4% in the random-window control. The vast majority (92%) of these events were associated with the IS6 family (**Figure 5c**). Although our sequence-based approach cannot confidently distinguish transposase-catalyzed cointegration from homologous recombination between pre-existing IS elements that serve only as regions of homology^44,45^, these results identify IS6-mediated fusion as a major and recurrent route to multireplicon plasmid formation.

## Discussion

From an evolutionary perspective, the abundance of multireplicon plasmids is puzzling. Multireplicon plasmids often carry large genetic cargo, potentially imposing substantial biological costs, as larger plasmids place greater demands on the DNA replication, transcription, and translation machinery. Consistent with this, plasmid costs scale with both plasmid length and the number of AMR genes they encode^46^. In addition, large fusion plasmids are more likely to carry redundant or conflicting regulatory systems^47,48^ and are more prone to genetic instability driven by recombination between homologous regions or other mobile genetic elements^31^.

Under these assumptions, multireplicon plasmids should be unstable in natural populations. However, this and other studies^14,20,31^ have recently shown that multireplicon plasmids are relatively abundant among sequenced plasmids. This mirrors the classical plasmid paradox^49^, which asks why plasmids are widespread despite the biological costs they impose on their hosts. Our work helps frame this apparent paradox by showing that multireplicon plasmids are enriched in predicted transfer-associated features and clinically relevant traits, including antimicrobial, metal and biocide resistance, and virulence genes, and exhibit broader observed host distributions (**Figures 1 and 2**). These properties may promote their persistence and dissemination, but they also raise the question of how multireplicon plasmids acquire and retain such functions.

Plasmid fusion followed by genomic rearrangements provides a compelling possibility^14,31^, at least for some multireplicon plasmids. By combining the gene repertoires of different plasmids, fusion can generate novel combinations of advantageous traits while establishing physical linkage between functions that would otherwise reside on separate molecules, potentially facilitating their inheritance and transmission. This process may be particularly effective because many of the traits examined here can be gained in a functionally dominant manner^50^: fusion with a plasmid bearing a resistance determinant or a complete transfer module may be sufficient to confer the corresponding resistance or mobility phenotype on the resulting multireplicon plasmid. Subsequent rearrangements could generate further structural variation, including variants that lose costly or unnecessary genetic cargo while retaining advantageous functions. Selection could then favor multireplicon configurations with more favorable combinations of traits. The abundance of multireplicon plasmids may therefore reflect both their recurrent formation and their differential persistence.

An additional potential benefit is that the combination of replicons in a multireplicon plasmid could facilitate persistence across different bacterial hosts^13,51^. This could help explain the broader observed host distributions of multireplicon plasmids in our dataset (**Figure 2**). Importantly, not every replication system needs to be functional in every host. Instead, one replication system could support maintenance of the entire multireplicon plasmid, allowing genes originally carried by another plasmid to persist even in hosts where its replication system would not function independently. Whether this mechanism contributes to the broader observed host distributions of multireplicon plasmids remains to be experimentally established.

Several of our findings are consistent with this evolutionary scenario. First, replicon pairs exhibited non-random associations (**Figure 3**), suggesting that certain combinations arise or persist more frequently than others. These patterns could reflect differences in the frequency or mechanisms of multireplicon formation, replicon compatibility, or selection. Second, some replicon pairs exhibited conserved spacing and high phylogenetic congruence, with the latter also positively associated with congruence between their flanking regions (**Figure 4, Supplementary Figure 12**). These patterns are consistent with stable physical linkage and a shared evolutionary history, although alternative explanations, including recent plasmid or host clonal expansion and recurrent recombination among closely related plasmids, cannot be excluded. In contrast, other replicon pairs showed little phylogenetic congruence, which could reflect multiple independent formation events or the erosion of a shared evolutionary history through recombination and replicon turnover. Together with the diverse genetic signatures identified at fusion junctions (**Figure 5**), these observations suggest that multireplicon plasmids arise through different molecular mechanisms and follow heterogeneous evolutionary trajectories.

Our work is not without limitations. Due to the intense global surveillance of AMR, genomic databases are heavily skewed toward clinically and epidemiologically relevant pathogens, particularly from *Enterobacterales* and *Firmicutes*. Although we removed highly similar plasmids to mitigate the effects of redundant sampling, our estimates of multireplicon plasmid abundance and their enrichment in mobility-associated features, antimicrobial, metal and biocide resistance genes, and virulence determinants may not reflect those of less intensively sampled, nonclinical microbes. This bias extends to replicon detection: 19,035 of the 72,556 PLSDB plasmids (26%) lacked a detectable replicon, highlighting the limitations of current replicon typing tools. Incomplete replicon detection may bias our analyses toward well-characterized taxa and replicon families, potentially underestimating the prevalence of multireplicon plasmids or misclassifying them as single-replicon plasmids. Similarly, the absence of particular replicons from a given taxon may reflect limited sequencing or annotation rather than true host-range boundaries. Moreover, because sequence databases contain only plasmids that persisted long enough to be sampled, the observed abundance of multireplicon plasmids and their observed properties reflects a combination of their formation, persistence, and clonal expansion. Finally, our analyses rely on software-based annotation of plasmid traits, including replicons, mobility, AMR genes, and virulence factors. Although these tools are standard in the field, experimental studies are needed to establish the functionality of these features and determine how they contribute to plasmid fitness, stability, and host range.

Despite these limitations, our data support a model in which multireplicon plasmids exist along a continuum from transient fusion intermediates to more stable genetic entities. Some likely represent short-lived intermediates in brief fusion-fission cycles, as recently proposed^14,31^, whereas others show structural and phylogenetic patterns compatible with long-term persistence. Multireplicon plasmids may therefore act as evolutionary testbeds, where unstable combinations are resolved or lost, while others persist. Overall, our findings establish multireplicon plasmids as widespread genetic platforms enriched in transfer associated sequence components and clinically relevant genes, while their non-random replicon associations, conserved spatial organization, phylogenetic congruence, and breakpoint genetic signatures provide testable hypotheses to investigate how these plasmids form and persist.

## Methods

### Dataset deduplication and metadata

First, all available plasmid sequences were downloaded from PLSDB^28^ v2024_05_31_v2. Plasmid-associated metadata was obtained from PLSDB, including replicon number and typing, mobility whereas host genus was extracted from the description field. Virulence factor information was extracted from VFDB^52^ for the “plasmid elements” table genes in PLSDB. Replicon annotations were obtained directly from the MOB-typer output distributed with PLSDB^53^. MOB-typer identifies replicon matches by BLASTn against the MOB-suite replicon database using its default thresholds: E-value ≤ 1 × 10^-5^, nucleotide identity ≥ 80%, query coverage ≥ 80%, and a minimum hit length of 80 bp. Fragmented alignments assigned to the same replicon–contig combination are collapsed, and overlapping records are filtered using a 5-bp overlap threshold, retaining the better-supported hit based on bit score. Plasmids were classified according to the number of replicon types reported by MOB-typer. Repeated hits assigned to the same replicon type at distinct, non-overlapping genomic loci were counted separately and could therefore result in classification as a multireplicon plasmid; this occurred in 307 of 23,925 representative plasmids (1.28%). Plasmids then were stratified by replicon count as defined by MOB-typer, and plasmids lacking replicons (n=19,035) were excluded.

To remove redundancy while preserving biological diversity, we performed a graph-based deduplication using pairwise distances computed with Mash^54^, which implements a *k*-mer–based MinHash approach to approximate sequence similarity. Distances were computed on predicted amino acid sequences using parameters (*k*-mer size = 9, sketch size = 2,000). First, all identical plasmids (mash distance = 0) were collapsed. Then, for each set of plasmids with the same number of replicons, an undirected similarity graph was constructed by connecting plasmid pairs with Mash distance ≤ 0.01, corresponding to near-identical genomic content. Community detection was then applied independently to each stratum using the Louvain algorithm^55^, yielding clusters of highly similar plasmids with the same number of replicons. For each cluster, the largest plasmid was selected as a single representative plasmid. Plasmids not connected to any other sequence were treated as singleton clusters and retained. A total of 23,925 out of 72,556 original plasmids were retained.

### Multivariable models of gene content

To assess whether associations between multireplicon status and functional gene presence persisted after adjustment for dataset covariates, we fitted separate binomial generalized linear models (logistic regressions with a logit link) for the presence or absence of at least one AMR, metal/biocide resistance, or virulence gene using the glm function from the statsmodels^56^ library in Python. Each plasmid contributed one binary response per functional category; additional gene copies did not affect this response. Multireplicon status was the focal explanatory variable. Models were adjusted for log10-transformed plasmid length, predicted mobility, host genus, and isolation source. Host genera represented by fewer than 200 plasmids were grouped as “Other”. Isolation source was classified as Human (n = 7,942), ND/Other (n = 5,777), Animal (n = 3,767), Food (n = 1,910), Soil (n = 1,771), Wastewater (n = 1,052), Plant (n = 854), or Water (n = 852). Adjusted odds ratios were obtained by exponentiating model coefficients, and 95% confidence intervals by exponentiating their corresponding confidence limits. Statistical significance was assessed using two-sided Wald tests, with Benjamini–Hochberg correction across the three functional outcomes.

### Length additivity analysis

To test how plasmid length scales with the number of replicons, we performed a genus-specific analysis to account for possible differences in length depending on the host. Within each genus, plasmids with exactly one replicon were used to define replicon-specific baseline lengths, computed as the median plasmid length per replicon. Only replicon–genus combinations supported by at least 10 single-replicon plasmids were retained. For each genus and each distinct multireplicon with sufficient support (at least five plasmids sharing the same replicons), the expected additive length was calculated as the sum of baseline medians.

### Mobility-associated features and predicted mobility classification

Mobility-associated sequence features and predicted mobility classes were analysed in 23,925 representative plasmids. MOB-typer annotations were used to analyse the presence of mobility associated features (relaxase, mating-pair formation (MPF) and origin of transfer), individually or in combination. We also used the MOB-typer categories based on the predicted presence of these components to classify plasmids as conjugative, mobilizable or non-mobilizable. To independently predict plasmid mobility, we used CONJScan protein predictions and oriT detection using OriTDB^57^. CONJScan analyses used MacSyFinder 2.1.6^58^, MacSyLib 1.0.4 and CONJScan^59^ 2.1.0, with the CONJScan/Plasmids models and ––db-type unordered. Accepted HMM hits had an independent E-value ≤0.001 and profile coverage ≥50%. The experimental OriTDB reference contained 116 unique queries. Queries were searched against plasmid sequences using BLASTn 2.16.0+ with –task blastn-short. Accepted alignments required an E-value <0.01, nucleotide identity >80% and query-span coverage >80%. Among overlapping hits, the hit with the lowest E-value was retained. Using the mobility framework described by Ares-Arroyo et al.^37^, plasmids were assigned to four mutually exclusive categories: pCONJ, with a complete MPF system and a relaxase; pMOB, with a relaxase but no complete MPF system; pOriT, with a detected oriT but no relaxase; and pNT, without a detected relaxase or oriT. Plasmids encoding at least two distinct structural MPF proteins without a complete MPF system were predicted to bear an incomplete conjugation apparatus (pdCONJ, n=2,072). Depending on whether they carried a relaxase, an oriT, or neither, these were classified into pMOB, pOriT, or pNT categories, respectively.

Associations with multireplicon status were evaluated using separate binomial logistic regression models for each feature, feature combination and mobility category. Models compared membership in each category with membership in all remaining categories. Models included multireplicon status, log10-transformed plasmid length, host genus and isolation source as variables. Host genera represented by fewer than 200 plasmids were grouped as “Other”. Adjusted odds ratios and 95% confidence intervals were obtained by exponentiating the multireplicon coefficient and its Wald confidence limits; two-sided Wald tests provided p-values. Benjamini–Hochberg correction was applied within separate testing families: the six MOB-typer feature endpoints together with predicted conjugative status, the three native MOB-typer mobility categories, and the four independent functional categories.

### Observed host distribution analysis

To identify replicons with a potential broader observed host distribution in multireplicon plasmids, we analyzed the full, non-deduplicated PLSDB dataset of 72,556 plasmids. Host genera were obtained from PLSDB taxonomic metadata, excluding non-informative annotations such as “uncultured”, “bacterium”, “metagenome”, or “environmental”. No replicon was detected in 19,035 plasmids (26.2%), whereas 37,525 (51.7%) were classified as single-replicon and 15,996 (22.0%) as multireplicon.

For each replicon, we first identified the genera in which it occurred in a single-replicon plasmid. We then checked whether it occurred in additional genera in multireplicon plasmids, counting each additional genus as a separate candidate case. To determine the taxonomic breadth of each case, we compared the additional genus/genera with all genera in which the replicon occurred in single-replicon plasmids and retained the lowest taxonomic level between the new genus and the ones in which it appeared as a single-replicon. This procedure identified 289 candidate cases: 187 at the genus level, 55 at the family level, 41 at the order level, two at the class level, and four at the phylum level.

We further assessed these cases using 1,863,797 bacterial assemblies from the 78 implicated genera in the AllTheBacteria database^39^. For each case, we searched for the replicon in assemblies from the additional genus. We discarded 66 cases in which it was detected as the only replicon in at least one assembly, even if it occurred with other replicons in other assemblies from that genus. Among the remaining cases, we considered a case supported when the replicon was detected in the same assembly with at least one of the other replicons present in the corresponding PLSDB multireplicon plasmid in that genus. Cases in which the replicon was not detected, or was detected only with replicons other than those found in the corresponding PLSDB plasmids, were considered not supported. Importantly, through this procedure only instances that are not supported are removed, but no new inferences of host range are created from AllTheBacteria database. This procedure externally supported 114 cases of the original 289 (39.4%). The results were visualized using a Sankey plot linking the taxa in which each replicon occurred in single-replicon plasmids to the additional genera in which it occurred in multireplicon plasmids. Each case was linked to all corresponding single-replicon host taxa, so a case could appear as several links connecting a baseline taxa with a replicon in the plot but was counted only once in the analysis.

### Replicon co-occurrence network

To characterize patterns of replicon co-occurrence on plasmids, we constructed a replicon co-occurrence network. Only plasmids from the full PLSDB database belonging to six predefined genera (*Escherichia*, *Klebsiella*, *Enterobacter*, *Salmonella*, *Staphylococcus*, and *Enterococcus*) were considered. Co-occurrence counts were computed for all pairs of replicons by counting the number of plasmids in which both replicons were present. These counts were used to construct a symmetric replicon–replicon co-occurrence matrix, from which an undirected weighted network was generated, with nodes representing replicon types and edge weights corresponding to absolute co-occurrence counts; replicons not co-occurring with any other replicon were removed. Community structure in the weighted network was inferred using the Louvain modularity optimization algorithm. Two large communities were identified by this procedure. To allow overlapping community membership, a soft assignment scheme was applied in which, for each replicon, weighted connectivity toward each of the two communities was computed as the sum of edge weights linking the replicon to members of the community, and replicons were assigned to a community if at least 25% of their total weighted connectivity to the two communities was directed toward that community.

### Cell-level co-occurrence analysis

To estimate cointegration frequency, that is, the tendency of a replicon pair to reside on a single multireplicon plasmid rather than two separate ones, we first grouped all plasmids belonging to the same assembly and then retrieved the replicon types detected on each plasmid. Only assemblies containing at least two distinct replicon types were considered. In total, 41,926 plasmids were retained. For every unordered pair of replicons (A,B) present within the same genome, we recorded whether both replicons were carried on the same plasmid or in different plasmids. Across all genomes, this yielded a binary outcome for each genome–replicon-pair instance, indicating same-plasmid versus different-plasmid localization. For each replicon pair, the cointegration frequency was defined as P(same plasmid∣same genome) computed as the proportion of genomes in which replicons A and B co-occurred on the same plasmid among all genomes where both replicons were present. Only replicon pairs observed together in at least five genomes were retained for downstream analysis. Statistical uncertainty in cointegration frequencies was quantified using exact binomial confidence intervals (95% confidence level) based on the Clopper–Pearson method^60^. Replicon pairs were then classified into three categories: strong cointegration, defined by a cointegration frequency ≥ 0.8 and a lower confidence bound ≥ 0.6; strong repulsion, defined by a cointegration frequency ≤ 0.2 and an upper confidence bound ≤ 0.4; and mixed, encompassing all remaining cases.

To assess whether cointegration frequencies were influenced by the overrepresentation of particular bacterial genera, we repeated the cell-level analysis using two complementary host-genus balancing approaches. First, frequencies were calculated separately within each genus. For each replicon pair, we then calculated an equal-weight genus mean by averaging the genus-specific estimates supported by at least five genomes. Second, we performed 300 subsampling iterations with representation balanced across host genera. In each iteration, up to 100 genomes were sampled without replacement from each genus, thereby limiting the contribution of highly represented genera while retaining all available genomes from genera represented by fewer than 100 genomes. Cointegration frequencies were recalculated in each balanced dataset, and the balanced estimate for each pair was defined as the mean across iterations in which that pair was observed. A fixed random seed was used. Agreement between the original and balanced estimates was assessed using Pearson and Spearman correlations.

### Replicon topological distance analysis

To quantify the relative genomic positioning of replicons within multireplicon plasmids, we analyzed the MOB-typer intermediate files from BLAST^61^ to identify the genomic coordinates of all assigned replicons. Replicon positions were defined using the start coordinate of the alignment on the plasmid sequence, and plasmid length was obtained directly from the BLAST-reported subject length. For each plasmid, all unordered pairs of assigned replicons were considered, and pairwise distances were computed assuming circular plasmid topology, defined as the minimum of the absolute coordinate difference and its complement with respect to plasmid length. Distances were reported both in absolute base pairs and normalized by plasmid length. To quantify how consistently each replicon pair maintained its spacing across plasmids, we calculated the coefficient of quartile variation (CQV) from the distribution of relative distances for each pair, defined as (Q3 − Q1)/(Q3 + Q1), where Q1 and Q3 are the first and third quartiles. CQV measures relative dispersion in a robust, scale-independent manner: low values indicate conserved spacing across plasmids, whereas high values indicate highly variable spacing. Because it is normalized by the magnitude of the distribution, CQV is appropriate for comparing variability among replicon pairs with substantially different typical distances, unlike raw variance or interquartile range, which increase with scale. To construct the null distributions, replicon positions were sampled independently from a uniform distribution along a circular plasmid, and the shortest normalized distance was calculated as for the observed data. For the CQV analysis, simulations preserved the number of observations for each replicon pair and were repeated 1,000 times using a fixed random seed. Replicon pairs represented in fewer than five plasmids were excluded from this analysis.

### Replicon phylogenetic congruence analysis

To assess evolutionary correspondence between co-occurring replicons, we compared the phylogenetic structure of each replicon across the set of deduplicated plasmids in which a given pair co-occurred. For each plasmid, replicon sequences were identified from MOB-typer outputs and extracted from the original plasmid sequences. Replicon nucleotide sequences were aligned independently with MUSCLE^62,63^ v5.3 with default parameters. Maximum-likelihood phylogenies were built in R using the phangorn package^64^. For each alignment, an initial tree was inferred by Neighbor-Joining based on maximum-likelihood distances (dist.ml), and subsequently optimized under the GTR substitution model with gamma-distributed rate heterogeneity using optim.pml (optGamma = TRUE, optInv = FALSE). For each replicon pair, patristic distance matrices were computed from the corresponding trees and restricted to plasmids represented in both phylogenies; pairs represented by fewer than four shared plasmids were excluded. Phylogenetic congruence between replicons was quantified using Mantel tests based on Pearson correlation between patristic distance matrices, with 9,999 permutations. In parallel, mirror-tree correlations were calculated from vectorized patristic distances, and significance was assessed by permuting tree tip labels 2,000 times to generate empirical null distributions. For selected pairs, phylogenetic congruence was visualized using midpoint-rooted tanglegrams. As a control, the same procedure was applied to the same replicon pairs when coexisting within the same assembly but located on separate single-replicon plasmids.

To determine whether phylogenetic congruence between co-occurring replicons was accompanied by conservation of their surrounding genomic regions, we examined the local sequence context of the replicon pairs included in the phylogenetic congruence analysis. Replicon coordinates and strand orientation were obtained from the MOB-typer BLAST results, and the corresponding regions were extracted from the original plasmid sequences. For each replicon, we extracted up to 1 kb on each side, excluding the replicon itself and accounting for circular plasmid topology. When two replicons were less than 2 kb apart, the sequence between them was split equally between their flanking regions to avoid overlap. The upstream and downstream flanks were concatenated and reverse-complemented as implemented for replicons on the negative strand. Each context sequence was represented as a set of distinct 15-mers. Pairwise sequence dissimilarity between plasmids was calculated using the Jaccard distance. For each replicon pair, we constructed two distance matrices using the same plasmids, one for each replicon’s flanking context. Congruence between the two context-distance matrices was assessed using a Pearson Mantel test with 99 permutations. Finally, the resulting flanking-context Mantel coefficients were compared with the corresponding replicon phylogenetic Mantel coefficients using Pearson and Spearman correlations across replicon pairs.

### NUCmer-based plasmid containment analysis

To identify plasmid containment relationships consistent with fusion events, we performed whole-sequence nucleotide alignments using the nucmer module from MUMmer v4^65^. Plasmids were stratified by their number of replicons and aligned against all plasmids with a higher number of replicons. Alignments were performed using the *--maxmatch* option to sensitively detect all maximal exact matches, and alignment coordinates were extracted using *show-coords*. For each alignment, plasmids were classified as smaller or larger based on their total length, and containment coverage was defined as the fraction of the smaller plasmid covered by the aligned sequence. Only alignments with at least 90% coverage of the smaller plasmid and at least 90% nucleotide identity were retained. Only cases exhibiting a single hit (one continuous sequence) were kept, excluding plasmids with internal duplications or complex mosaic architectures, also, cases where the small plasmid carried more replicons were excluded.

### Analysis of cointegration breakpoints

To identify putative homologous sequences present at the two breakpoints of inferred plasmid fusion events, we used the corresponding NUCmer alignment file to recover the coordinates of the two breakpoints as the start and end positions of that interval. Around each breakpoint, we extracted a sequence window of ±500 bp (1 kb total) from the original plasmid. Then, the two breakpoint-centered windows were compared against each other using BLASTn^61^ with dust masking disabled, soft masking disabled, and a minimum nucleotide identity threshold of 95%. When multiple local alignments were detected between the two windows, only the single best hit per breakpoint pair was retained, ranked by alignment length. A value of 0 bp was assigned when no alignment met the search criteria. Homologous sequences were then clustered across events using cd-hit-est v4.8.1^66^ with a sequence identity threshold of 95%, and default coverage parameters. To assess whether breakpoint-associated homology exceeded that expected by chance, we generated a matched random control dataset using the same plasmid set and the same number of breakpoint pairs. For each fusion event, two random positions were sampled on the same plasmid, and ±500 bp windows were extracted around these random positions using the same procedure as for the breakpoint data. These random window pairs were compared with BLASTn using identical parameters.

To assess IS involvement in plasmid fusion events, we analyzed breakpoint-centered sequence windows around fusion breakpoints. For each breakpoint, we widened the extracted sequence windows to ±2000 bp centered on the breakpoint (4 kb total), and used digIS v1.2^67^ to identify IS elements using default parameters. To avoid counting the same IS-associated region twice when windows overlapped, we required matching IS annotations to correspond to two distinct, nonoverlapping genomic regions. When more than 1 IS was detected in the window, the closest one to the breakpoint was selected. Breakpoint pairs corresponding to each fusion event were then classified based on IS presence at the two breakpoints as “0 IS”, “1 IS”, “2 IS (duplicated)”, or “2 IS (different)”, depending on whether IS elements were absent, present at one breakpoint, or present at both breakpoints with identical or distinct annotations. It is important to note that the homology-based and IS-based analyses detect different, potentially overlapping sequence signatures and should not be interpreted as mutually exclusive evidence for distinct fusion mechanisms.

## Author Contributions

I.d.Q., P.R.-M., R.S.R analysed the data and created the figures. V.F.L., C.H.R., and J.R.-B. provided technical support and conceptual advice. I.d.Q., V.F.L., and J.R.-B. conceived the project. V.F.L., and J.R.-B. supervised the project. All authors discussed and provided critical feedback during the analysis of the results. All authors wrote, edited, and reviewed the manuscript.

## Materials & Correspondence

Correspondence and requests for materials should be addressed to Val F. Lanza and Jerónimo Rodríguez-Beltrán.

## Data availability

Datasets generated and/or analyzed during the current study are included in the Source Data file with this paper and can be downloaded from the following repository https://github.com/nachodeq/Multireplicon_plasmids_paper.

## Code availability

The source code used to run the analyses and produce the results presented in this manuscript is available at https://github.com/nachodeq/Multireplicon_plasmids_paper.

## Acknowledgements

We thank Javier DelaFuente, Laura Álvaro, and João Gama for their helpful suggestions. Work in the evodynamics lab (https://evodynamicslab.com/) is supported by project PI23/01945, funded by the Carlos III Health Institute (ISCIII) and co-funded by the European Union; CIBER-Consorcio Centro de Investigación Biomédica en Red-(CIBERINFEC) CB21/13/00084; Convocatoria SEIMC-Fundación Soria Melguizo de Investigación 2021; and funded by the European Union (ERC, HorizonGT, 101077809). Views and opinions expressed are, however, those of the author (s) only and do not necessarily reflect those of the European Union or the European Research Council Executive Agency. Neither the European Union nor the granting authority can be held responsible for them. This work was supported by project PID2025-176358NB-I00 funded by MICIU/AEI/10.13039/501100011033 and FEDER, UE. IdQ is a recipient of “Ayudas Fundación Ramón Areces para la realización de Tesis Doctorales en Ciencias de la Vida y de la Materia 2025” grant. The project that gave rise to these results received the support of a fellowship from Fundación Ramón Areces. PRM is a recipient of a predoctoral PFIS grant (grant no. FI22/00265) from the Carlos III Health Institute (ISCIII), through the Recovery, Transformation and Resilience Plan and Next Generation EU from the European Union. RSR received a fellowship from Programa de Doutorado Sanduíche no Exterior (PDSE) of Coordenação de Aperfeiçoamento de Pessoal de Nível Superior (CAPES) [Grant Number 88881.128025/2025-01]. V.F.L. acknowledges support by a Miguel Servet contract from the Carlos III Health Institute (ISCIII) (grant no. CP22/00164), co-founded by the European Social Fund, ‘Investing in your future’.

## References

1. Murray, C. J. L. et al. Global burden of bacterial antimicrobial resistance in 2019: a systematic analysis. The Lancet 399, 629–655 (2022).

2. Tokuda, M. & Shintani, M. Microbial evolution through horizontal gene transfer by mobile genetic elements. Microb. Biotechnol. 17, e14408 (2024).

3. McInnes, R. S., McCallum, G. E., Lamberte, L. E. & Van Schaik, W. Horizontal transfer of antibiotic resistance genes in the human gut microbiome. Curr. Opin. Microbiol. 53, 35–43 (2020).

4. Castañeda-Barba, S., Top, E. M. & Stalder, T. Plasmids, a molecular cornerstone of antimicrobial resistance in the One Health era. Nat. Rev. Microbiol. 22, 18–32 (2024).

5. Rodríguez-Beltrán, J., DelaFuente, J., León-Sampedro, R., MacLean, R. C. & San Millán, Á. Beyond horizontal gene transfer: the role of plasmids in bacterial evolution. Nat. Rev. Microbiol. 1–13 (2021) doi:10.1038/s41579-020-00497-1.

6. Akhtar, P. & Khan, S. A. Two independent replicons can support replication of the anthrax toxin-encoding plasmid pXO1 of Bacillus anthracis. Plasmid 67, 111–117 (2012).

7. Banerjee, S. K., Luck, B. T., Kim, H. Y. & Iyer, V. N. Three clustered origins of replication in a promiscuous-plasmid replicon and their differential use in a PolA+ strain and a delta PolA strain of Escherichia coli K-12. J. Bacteriol. 174, 8139–8143 (1992).

8. Imanaka, T., Ano, T., Fujii, M. & Aiba, S. Two replication determinants of an antibiotic-resistance plasmid, pTB19, from a thermophilic bacillus. J. Gen. Microbiol. 130, 1399–1408 (1984).

9. Lee, J.-H. & O’Sullivan, D. J. Sequence Analysis of Two Cryptic Plasmids from *Bifidobacterium longum* DJO10A and Construction of a Shuttle Cloning Vector. Appl. Environ. Microbiol. 72, 527–535 (2006).

10. Oskam, L., Hillenga, D. J., Venema, G. & Bron, S. The large Bacillus plasmid pTB19 contains two integrated rolling-circle plasmids carrying mobilization functions. Plasmid 26, 30–39 (1991).

11. Oskam, L., Venema, G. & Bron, S. Plasmid pTB913 derivatives are segregationally stable in Bacillus subtilis at elevated temperatures. Plasmid 28, 70–79 (1992).

12. van Kranenburg, R. & de Vos, W. M. Characterization of multiple regions involved in replication and mobilization of plasmid pNZ4000 coding for exopolysaccharide production in Lactococcus lactis. J. Bacteriol. 180, 5285–5290 (1998).

13. Rozwandowicz, M. et al. Plasmids carrying antimicrobial resistance genes in Enterobacteriaceae. J. Antimicrob. Chemother. 73, 1121–1137 (2018).

14. Cazares, A. et al. Pre– and postantibiotic epoch: The historical spread of antimicrobial resistance. Science 390, eadr1522 (2025).

15. Cameranesi, M. M., Morán-Barrio, J., Limansky, A. S., Repizo, G. D. & Viale, A. M. Site-Specific Recombination at XerC/D Sites Mediates the Formation and Resolution of Plasmid Co-integrates Carrying a blaOXA-58– and TnaphA6-Resistance Module in Acinetobacter baumannii. Front. Microbiol. 9, 66 (2018).

16. Chavda, K. D. et al. Complete Sequence of a *bla*_KPC_-Harboring Cointegrate Plasmid Isolated from Escherichia coli. Antimicrob. Agents Chemother. 59, 2956–2959 (2015).

17. Chen, L. et al. Carbapenemase-producing Klebsiella pneumoniae: molecular and genetic decoding. Trends Microbiol. 22, 686–696 (2014).

18. Desmet, S. et al. Antibiotic Resistance Plasmids Cointegrated into a Megaplasmid Harboring the *bla*_OXA-427_ Carbapenemase Gene. Antimicrob. Agents Chemother. 62, e01448–17 (2018).

19. Rodrigues, C. et al. KPC-3-Producing Klebsiella pneumoniae in Portugal Linked to Previously Circulating Non-CG258 Lineages and Uncommon Genetic Platforms (Tn4401d-IncFIA and Tn4401d-IncN). Front. Microbiol. 7, (2016).

20. De Souza, H. C. A., De Oliveira Almeida, A. C., ConteJunior, C. A. & Panzenhagen, P. Multi-replicon Architecture Drives the Global Accumulation of Resistance to Antimicrobials, Biocides, and Metals in IncF and IncH Plasmids. Curr. Microbiol. 83, 241 (2026).

21. Peterson, B. C., Hashimoto, H. & Rownd, R. H. Cointegrate formation between homologous plasmids in Escherichia coli. J. Bacteriol. 151, 1086–1094 (1982).

22. Hua, X. et al. Cointegration as a mechanism for the evolution of a KPC-producing multidrug resistance plasmid in Proteus mirabilis. Emerg. Microbes Infect. 9, 1206–1218 (2020).

23. He, S. et al. Insertion Sequence IS26 Reorganizes Plasmids in Clinically Isolated Multidrug-Resistant Bacteria by Replicative Transposition. mBio 6, e00762 (2015).

24. Siguier, P., Filée, J. & Chandler, M. Insertion sequences in prokaryotic genomes. Curr. Opin. Microbiol. 9, 526–531 (2006).

25. Liu, Z., Tang, Y., He, M. & Xu, C. Molecular drivers of fusion plasmid: mechanistic insights and evolutionary implications. J. Antimicrob. Chemother. 80, 2902–2911 (2025).

26. Harmer, C. J. & Hall, R. M. Targeted Conservative Cointegrate Formation Mediated by IS26 Family Members Requires Sequence Identity at the Reacting End. mSphere 6, e01321–20 (2021).

27. Novick, R. P., Iordanescu, S., Surdeanu, M. & Edelman, I. Transduction-related cointegrate formation between staphylococcal plasmids: A new type of site-specific recombination. Plasmid 6, 159–172 (1981).

28. Molano, L.-A. G., Hirsch, P., Hannig, M., Müller, R. & Keller, A. The PLSDB 2025 update: enhanced annotations and improved functionality for comprehensive plasmid research. Nucleic Acids Res. 53, D189–D196 (2025).

29. Galata, V., Fehlmann, T., Backes, C. & Keller, A. PLSDB: a resource of complete bacterial plasmids. Nucleic Acids Res. 47, D195–D202 (2019).

30. Schmartz, G. P. et al. PLSDB: advancing a comprehensive database of bacterial plasmids. Nucleic Acids Res. 50, D273–D278 (2022).

31. Ipoutcha, T., Wang, Y., Rocha, E. P. C. & Penadés, J. R. Mobile genetic elements drive a plasmid fusion and deletion lifecycle shaping evolution and antimicrobial resistance. Nat. Commun. 17, 10849 (2026).

32. Smillie, C., Garcillan-Barcia, M. P., Francia, M. V., Rocha, E. P. C. & de la Cruz, F. Mobility of Plasmids. Microbiol. Mol. Biol. Rev. 74, 434–452 (2010).

33. Russell, A. D. Plasmids and bacterial resistance to biocides. J. Appl. Microbiol. 83, 155–165 (1997).

34. Pilla, G. & Tang, C. M. Going around in circles: virulence plasmids in enteric pathogens. Nat. Rev. Microbiol. 16, 484–495 (2018).

35. de Souza, H. C. A. et al. Understanding the Association of Plasmid Incompatibility Groups With Variable Antimicrobial Resistance Genotypes in Bacteria. MicrobiologyOpen 14, e70187 (2025).

36. Coluzzi, C., Garcillán-Barcia, M. P., De La Cruz, F. & Rocha, E. P. C. Evolution of Plasmid Mobility: Origin and Fate of Conjugative and Nonconjugative Plasmids. Mol. Biol. Evol. 39, msac115 (2022).

37. Ares-Arroyo, M., Coluzzi, C. & Rocha, E. P. C. Origins of transfer establish networks of functional dependencies for plasmid transfer by conjugation. Nucleic Acids Res. 51, 3001–3016 (2023).

38. Ares-Arroyo, M., Coluzzi, C., Moura De Sousa, J. A. & Rocha, E. P. C. Hijackers, hitchhikers, or co-drivers? The mysteries of mobilizable genetic elements. PLOS Biol. 22, e3002796 (2024).

39. Hunt, M. et al. AllTheBacteria: a community resource empowers biology and discovers novel peptide antibiotics. Preprint at 10.1101/2024.03.08.584059 (2024).

40. Mantel, N. The detection of disease clustering and a generalized regression approach. Cancer Research, 27(2 Part 1):209–220, 1967. (1967).

41. Hommola, K., Smith, J. E., Qiu, Y. & Gilks, W. R. A Permutation Test of Host-Parasite Cospeciation. Mol. Biol. Evol. 26, 1457–1468 (2009).

42. Varani, A., He, S., Siguier, P., Ross, K. & Chandler, M. The IS6 family, a clinically important group of insertion sequences including IS26. Mob. DNA 12, 11 (2021).

43. Watt, V. M., Ingles, C. J., Urdea, M. S. & Rutter, W. J. Homology requirements for recombination in Escherichia coli. Proc. Natl. Acad. Sci. 82, 4768–4772 (1985).

44. Harmer, C. J. & Hall, R. M. IS26 Family Members IS257 and IS1216 Also Form Cointegrates by Copy-In and Targeted Conservative Routes. mSphere 5, e00811–19 (2020).

45. Harmer, C. J. & Hall, R. M. IS26 and the IS26 family: versatile resistance gene movers and genome reorganizers. Microbiol. Mol. Biol. Rev. MMBR 88, e0011922 (2024).

46. Vanacker, M., Lenuzza, N. & Rasigade, J.-P. The fitness cost of horizontally transferred and mutational antimicrobial resistance in Escherichia coli. Front. Microbiol. 14, (2023).

47. Liu, Z. et al. Adaptive evolution of plasmid and chromosome contributes to the fitness of a blaNDM-bearing cointegrate plasmid in Escherichia coli. ISME J. 18, wrae037 (2024).

48. San Millan, A. & MacLean, R. C. Fitness Costs of Plasmids: a Limit to Plasmid Transmission. Microbiol. Spectr. 5, (2017).

49. Brockhurst, M. A. & Harrison, E. Ecological and evolutionary solutions to the plasmid paradox. Trends Microbiol. 30, 534–543 (2022).

50. Rodríguez-Beltrán, J. et al. Genetic dominance governs the evolution and spread of mobile genetic elements in bacteria. Proc. Natl. Acad. Sci. 117, 15755–15762 (2020).

51. Osborn, A. M., Da Silva Tatley, F. M., Steyn, L. M., Pickup, R. W. & Saunders, J. R. Mosaic plasmids and mosaic replicons: evolutionary lessons from the analysis of genetic diversity in IncFII-related replicons. Microbiology 146, 2267–2275 (2000).

52. Liu, B., Zheng, D., Zhou, S., Chen, L. & Yang, J. VFDB 2022: a general classification scheme for bacterial virulence factors. Nucleic Acids Res. 50, D912–D917 (2022).

53. Robertson, J. & Nash, J. H. E. MOB-suite: software tools for clustering, reconstruction and typing of plasmids from draft assemblies. *Microb*. Genomics 4, e000206 (2018).

54. Ondov, B. D. et al. Mash: fast genome and metagenome distance estimation using MinHash. Genome Biol. 17, 132 (2016).

55. Blondel, V. D., Guillaume, J.-L., Lambiotte, R. & Lefebvre, E. Fast unfolding of communities in large networks. J. Stat. Mech. Theory Exp. 2008, P10008 (2008).

56. Seabold, S. & Perktold, J. Statsmodels: Econometric and Statistical Modeling with Python. in 92–96 (Austin, Texas, 2010). doi:10.25080/Majora-92bf1922-011.

57. Liu, G. et al. oriTDB: a database of the origin-of-transfer regions of bacterial mobile genetic elements. Nucleic Acids Res. 53, D163–D168 (2025).

58. Néron, B., et al. MacSyFinder v2: Improved modelling and search engine to identify molecular systems in genomes. Peer Community J. 3, e28 (2023).

59. Cury, J., Touchon, M. & Rocha, E. P. C. Integrative and conjugative elements and their hosts: composition, distribution and organization. Nucleic Acids Res. 45, 8943–8956 (2017).

60. Clopper, C. J. & Pearson, E. S. THE USE OF CONFIDENCE OR FIDUCIAL LIMITS ILLUSTRATED IN THE CASE OF THE BINOMIAL. Biometrika 26, 404–413 (1934).

61. Ye, J., McGinnis, S. & Madden, T. L. BLAST: improvements for better sequence analysis. Nucleic Acids Res. 34, W6–W9 (2006).

62. Edgar, R. C. MUSCLE: a multiple sequence alignment method with reduced time and space complexity. BMC Bioinformatics 5, 113 (2004).

63. Edgar, R. C. Muscle5: High-accuracy alignment ensembles enable unbiased assessments of sequence homology and phylogeny. Nat. Commun. 13, 6968 (2022).

64. Schliep, K. P. phangorn: phylogenetic analysis in R. Bioinformatics 27, 592–593 (2011).

65. Marçais, G. et al. MUMmer4: A fast and versatile genome alignment system. PLOS Comput. Biol. 14, e1005944 (2018).

66. Li, W. & Godzik, A. Cd-hit: a fast program for clustering and comparing large sets of protein or nucleotide sequences. Bioinformatics 22, 1658–1659 (2006).

67. Puterová, J. & Martínek, T. digIS: towards detecting distant and putative novel insertion sequence elements in prokaryotic genomes. BMC Bioinformatics 22, 258 (2021).

