## Supplementary_Material for "Multireplicon plasmids show broad host distributions, high transfer potential, and are enriched in antimicrobial resistance genes"

1. Microbiology Department. Hospital Universitario Ramón y Cajal-IRYCIS, Madrid, Spain.
2. Escuela de Doctorado. Universidad Autónoma de Madrid.
3. Department of Clinical Analyses, Toxicology and Food Science, School of Pharmaceutical Sciences of Ribeirão Preto, University of São Paulo, Ribeirão Preto, São Paulo, Brazil.
4. Centro de Investigación Biomédica en Red de Enfermedades Infecciosas (CIBERINFEC), Instituto de Salud Carlos III, Madrid, Spain.
5. Translational Genomics and Bioinformatics Unit. IRYCIS. Madrid, Spain.

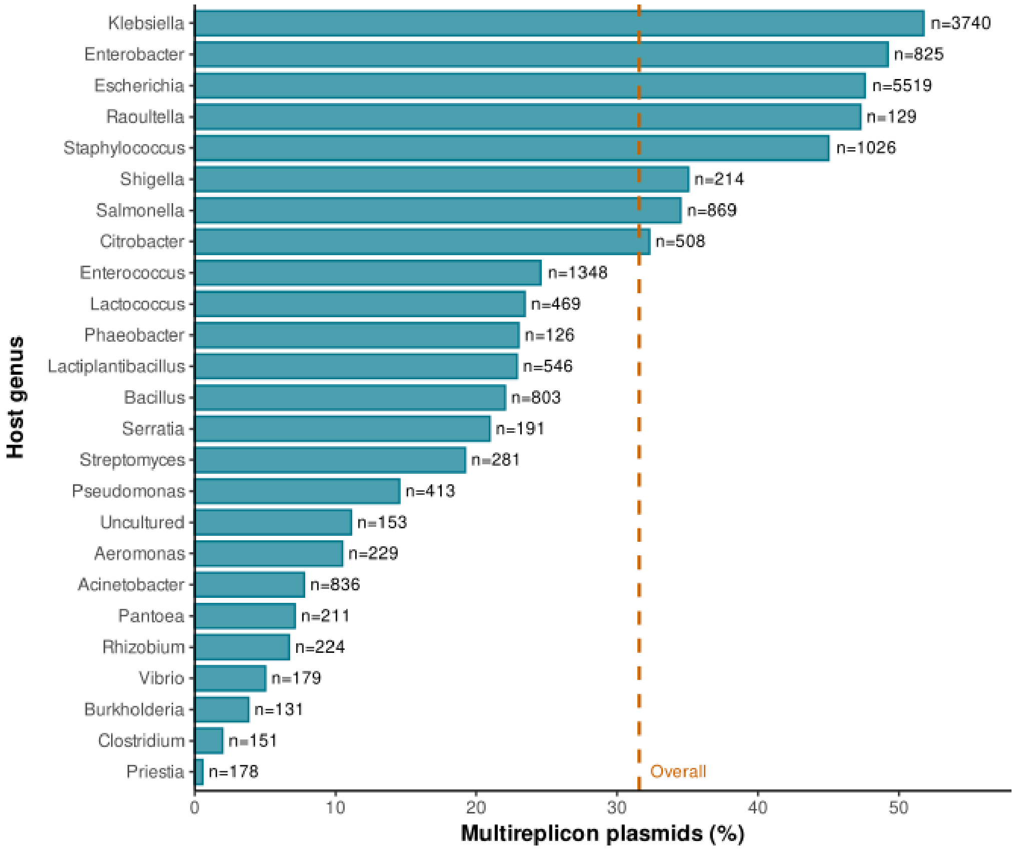


***Supplementary Figure 1. Multireplicon abundance and proportion across bacterial genera.*** *Bar plots show the N and % of multireplicon plasmids in the top 25 most abundant bacterial genera in the database*

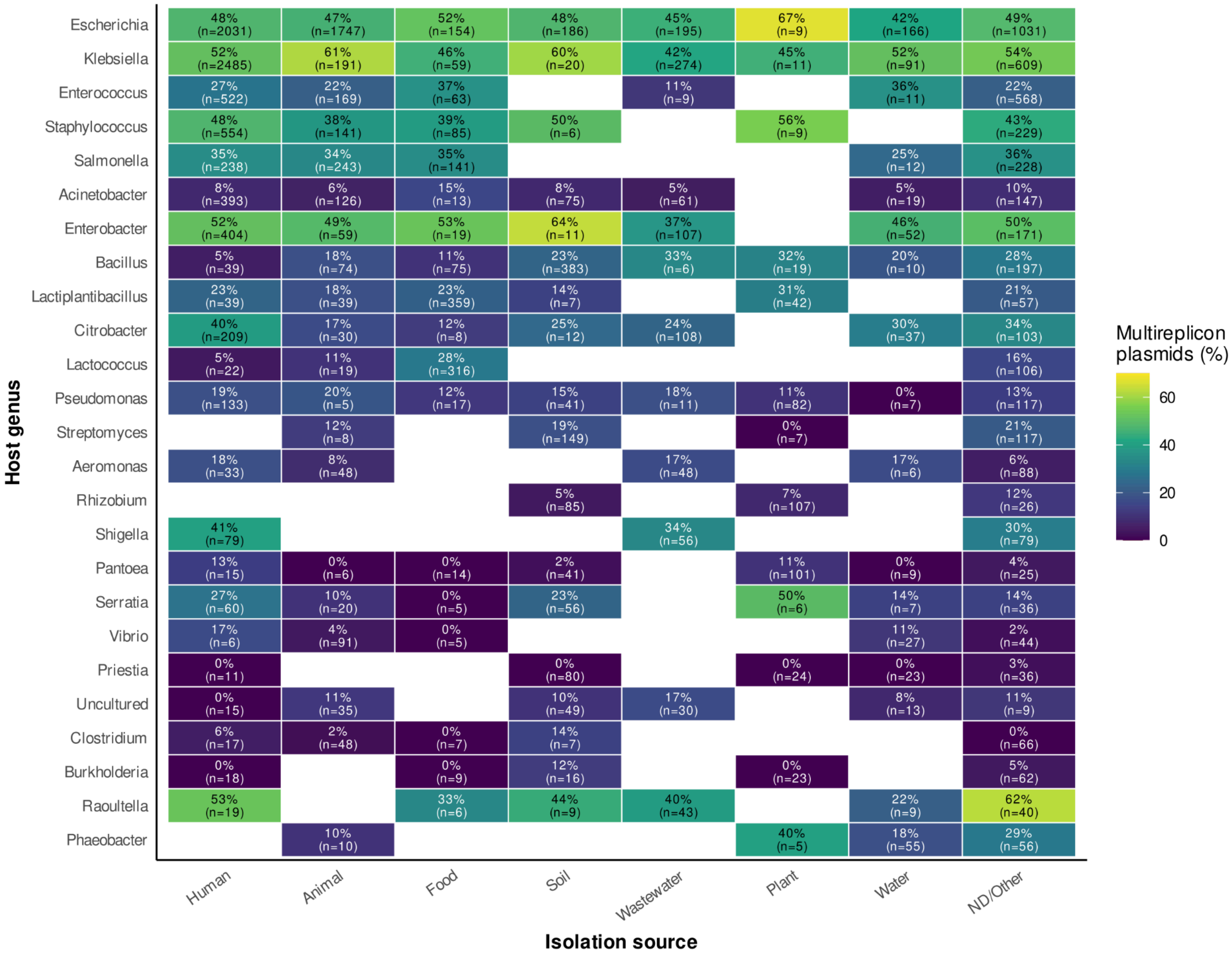


***Supplementary Figure 2. Isolation source and multireplicon prevalence.*** *Heatmap showing the proportion of multireplicon plasmids across isolation sources within the 25 most abundant host genera in the representative dataset. Each cell reports the percentage of multireplicon plasmids and the total number of plasmids in the corresponding genus-source combination. Blank cells indicate combinations with less than 5 observations.*

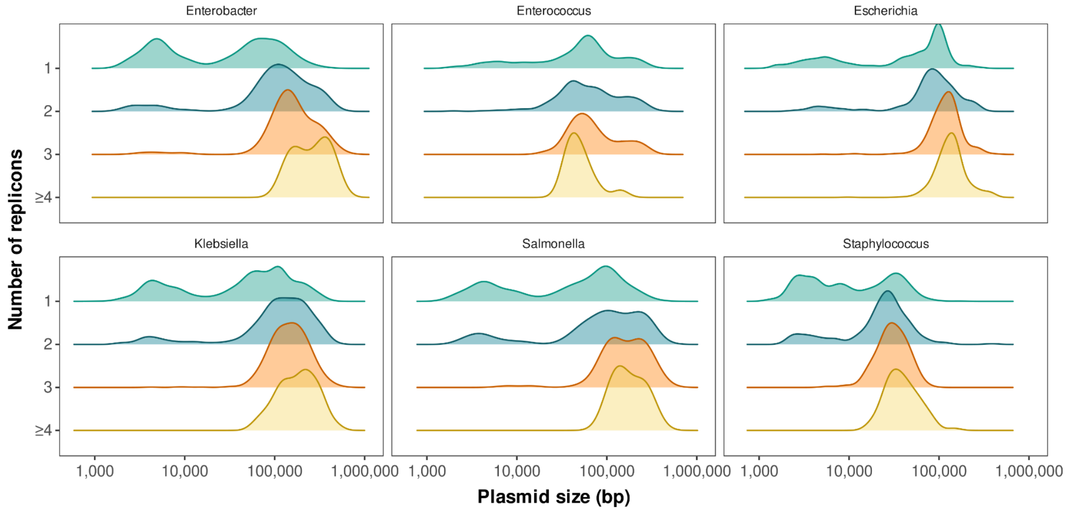


***Supplementary Figure 3. Plasmid length distributions stratified by replicon number across most represented bacterial genera in the dataset.*** *Kernel density estimates of plasmid sizes (bp, log scale) are shown for different numbers of replicons within each genus. Colors indicate the number of replicons per plasmid. Across all genera, plasmids with multiple replicons are consistently skewed toward larger lengths.*

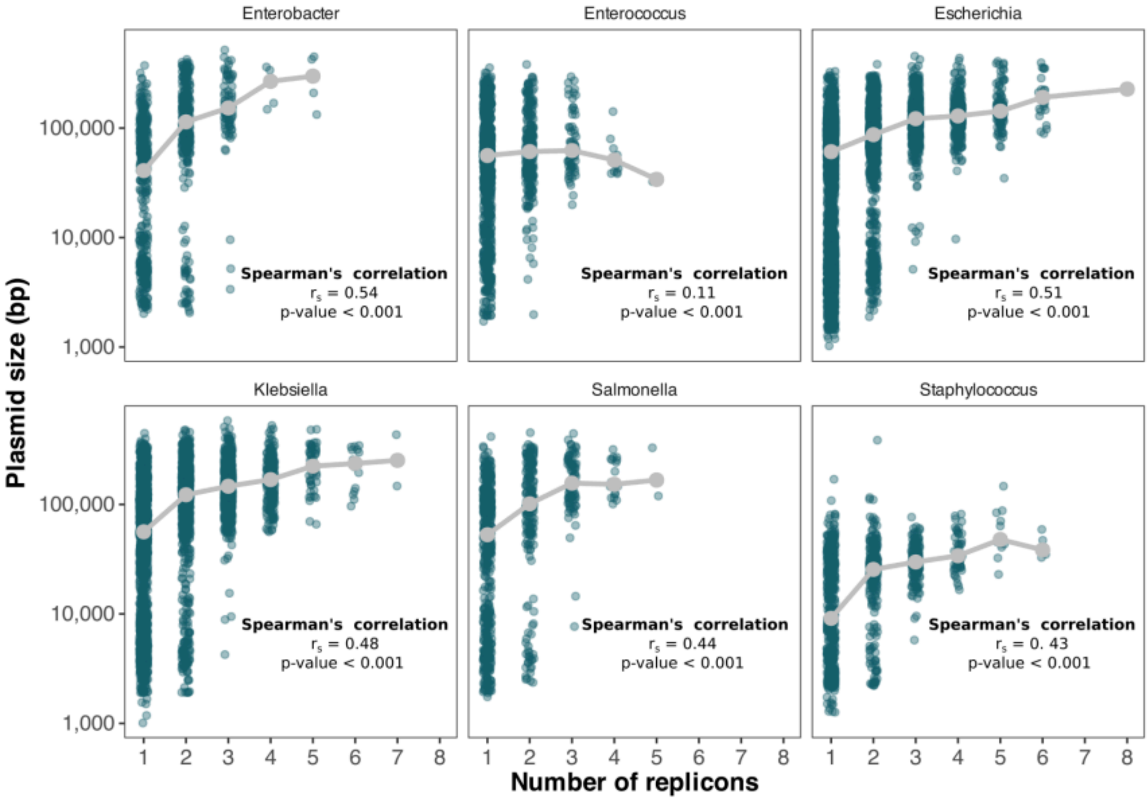


***Supplementary Figure 4. Relationship between plasmid size and replicon number across the most represented bacterial genera in the dataset.*** *Plasmid size as a function of replicon number across genera (each point represents a plasmid; y-axis on logarithmic scale); grey points and lines summarize the central trend, and correlation coefficients (Spearman's* $\rho$ and $p$*) are shown within each panel.*

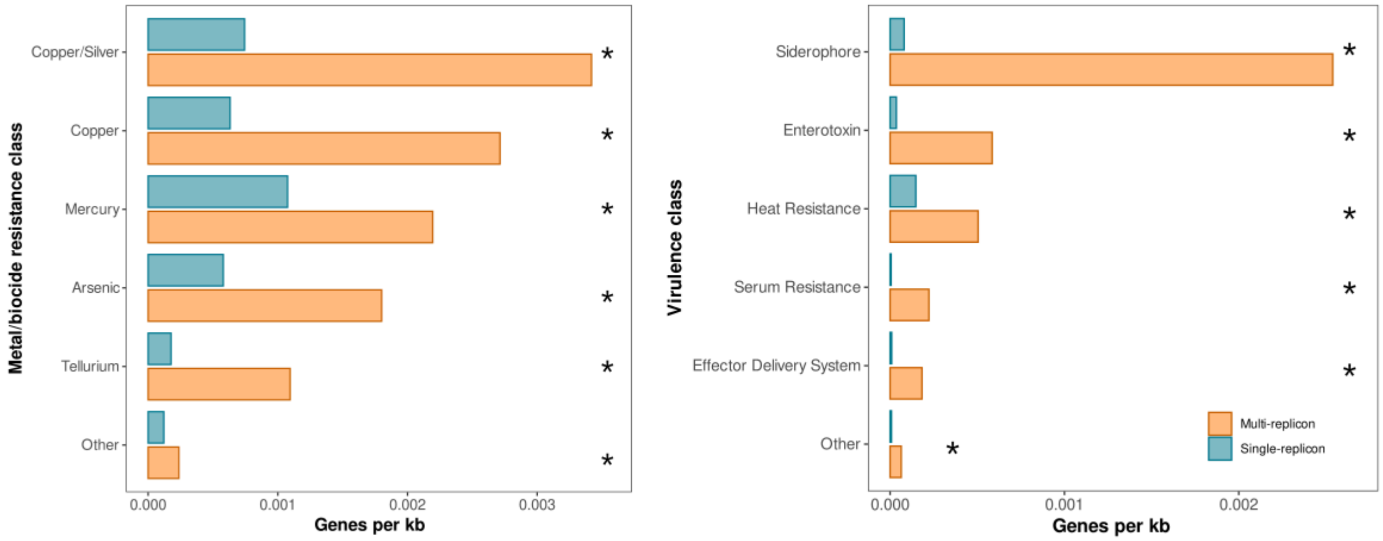


***Supplementary Figure 5. Enrichment of metal/biocide resistance and virulence gene classes in multi-replicon plasmids.*** *Bar plots show the mean gene density (genes per kb) for the most prevalent functional classes in single-replicon (blue) and multi-replicon (orange) plasmids. For each category, the top five classes ranked by abundance in multi-replicon plasmids are displayed, with remaining classes grouped as "Other." Across both metal/biocide resistance (left) and virulence (right) categories, multi-replicon plasmids consistently exhibit higher gene densities than single-replicon plasmids. Statistical significance was assessed using two-sided Mann–Whitney U tests with false discovery rate (FDR) correction; asterisks indicate significant differences (FDR < 0.001).*

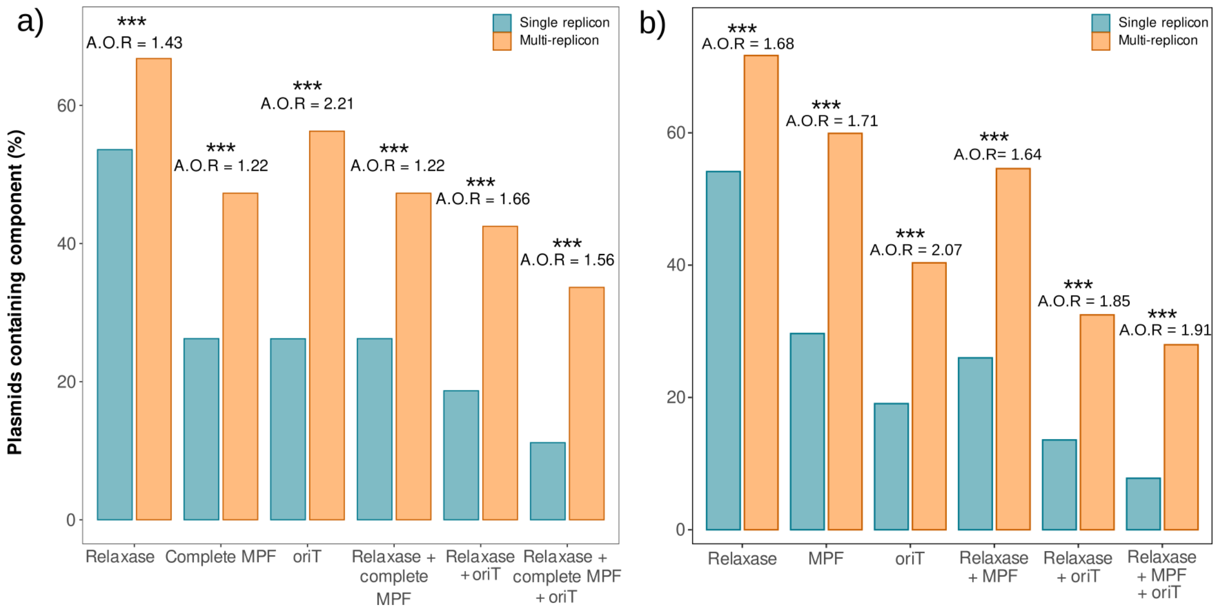


***Supplementary Figure 6. Predicted mobility-associated features in single- and multireplicon plasmids.*** *Predicted presence of relaxase, mating-pair formation (MPF) machinery, and origin-of-transfer (oriT), individually and in the indicated combinations by **a)** ConjScan and **b)** MOB-typer. Adjusted odds ratios (A.O.R.) compare multireplicon with single-replicon plasmids and were estimated using separate binomial logistic regression models adjusted for plasmid length, host genus, and isolation source.*

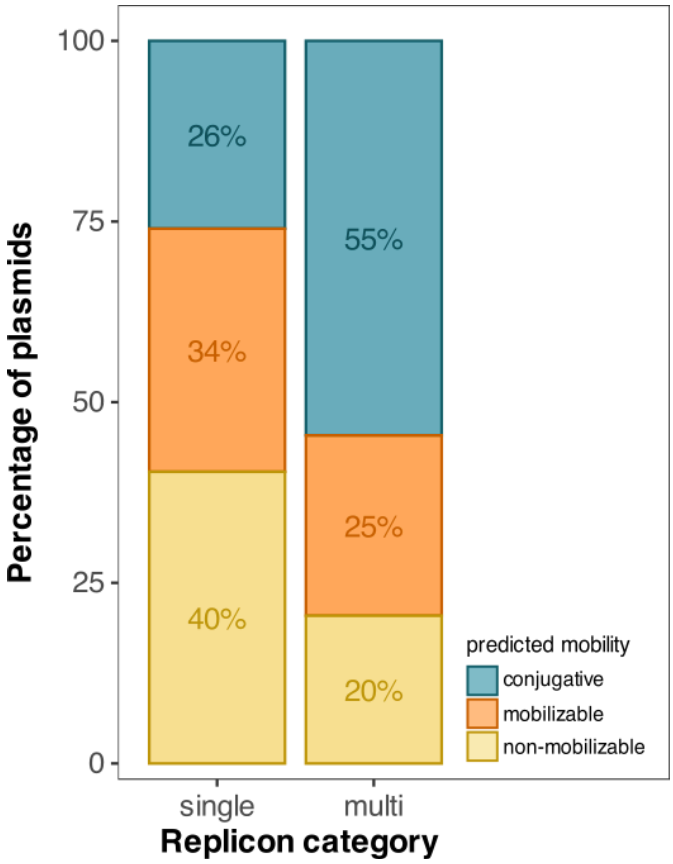


***Supplementary Figure 7. Predicted mobility classes of single-replicon and multireplicon plasmids using MOB-typer****. Stacked bars show MOB-typer classifications for single-replicon (n = 16,376) and multireplicon (n = 7,549) plasmids: conjugative (25.98% vs. 54.59%), mobilizable (33.65% vs. 24.96%), and non-mobilizable (40.37% vs. 20.45%). Binomial logistic regressions comparing each class with the remaining classes, adjusted for log10 plasmid length, host genus, and isolation source, yielded respective adjusted odds ratios of 1.64 (95% CI, 1.52–1.76), 1.11 (1.03–1.20), and 0.50 (0.47–0.54) for multireplicon versus single-replicon plasmids.*

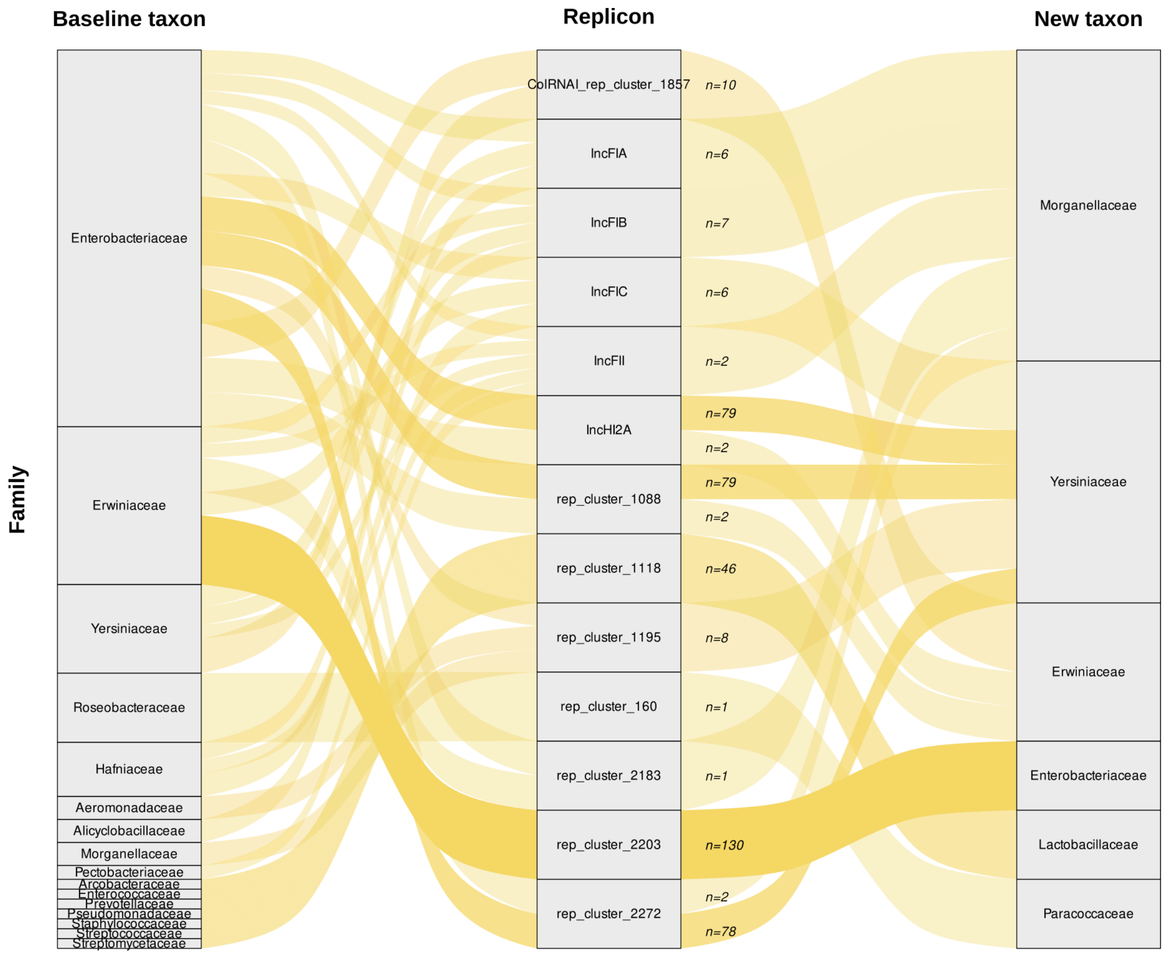


***Supplementary Figure 8. Replicons showing broader host distributions at the family level.*** *Baseline host families in which each focal replicon was detected on single-replicon plasmids are shown on the left, focal replicons in the center, and the additional host families in which they were observed on multireplicon plasmids on the right. Labels denoted indicate the number of assemblies supporting each event. Because one event may link to multiple baseline families at its minimum taxonomic-distance rank, it can contribute multiple visualization edges but is counted only once.*

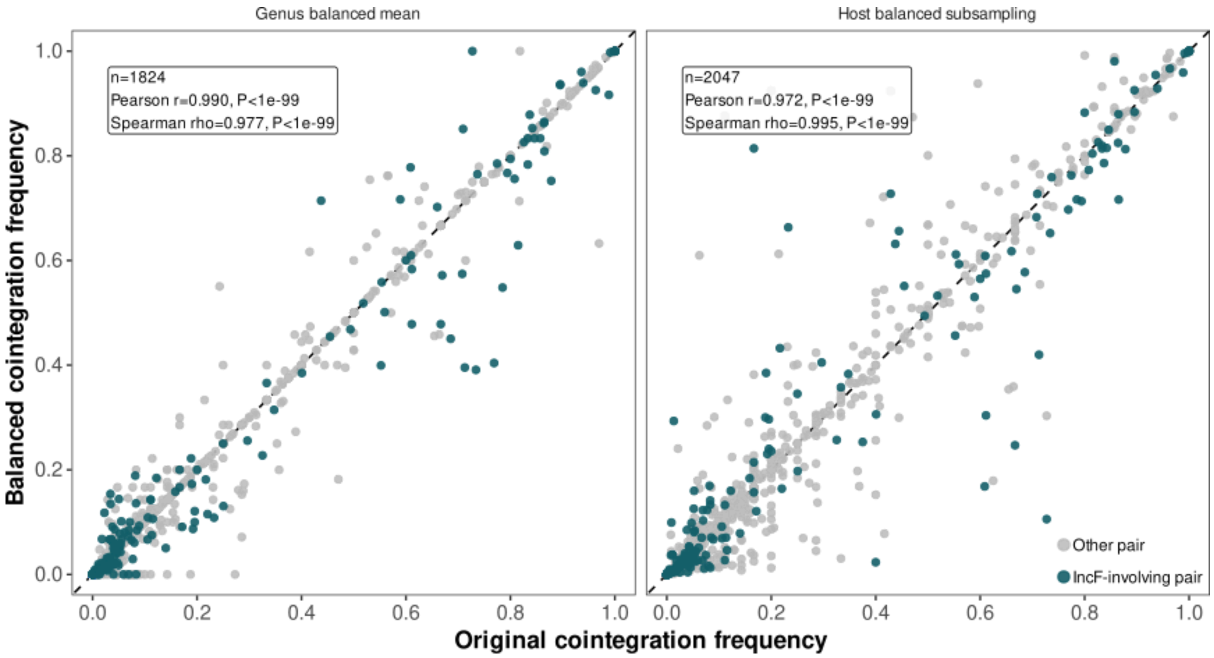


***Supplementary Figure 9. Comparison of cointegration frequencies before and after balancing host genera.*** *Each point represents a replicon pair observed together within the same genome in at least five genomes. The x-axis shows the cointegration frequency calculated from the complete dataset.* ***a)*** *Comparison with the equal-weight genus mean, calculated by averaging genus-specific frequencies supported by at least five genomes.* ***b)*** *Comparison with the mean frequency obtained across 300 subsampling iterations with representation balanced across host genera, each retaining up to 100 genomes per genus. To better reflect the effect of abundant replicons in the dataset, replicon pairs containing at least one IncF-family replicon are highlighted in green, whereas all other pairs are shown in grey. The dashed diagonal indicates equality between the original and balanced estimates.*

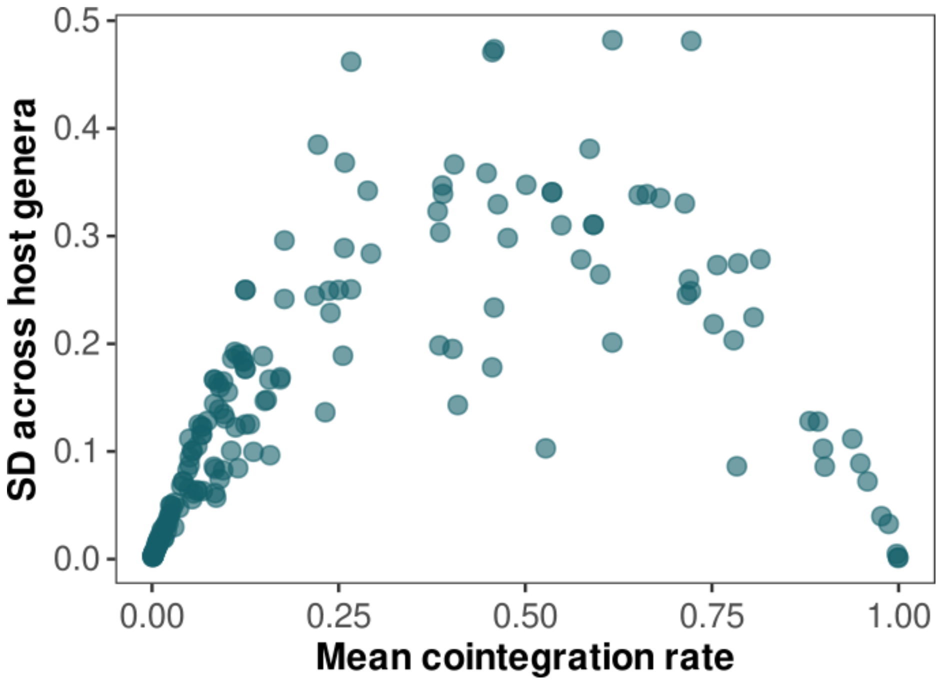


***Supplementary Figure 10. Variability of cointegration frequencies across host genera.*** *Each point represents a pair of replicons. The x-axis shows the mean cointegration frequency, while the y-axis indicates the standard deviation (SD) of this rate across genera. Replicon pairs with high or low cointegration frequencies (<0.2 or >0.8) tend to exhibit low variability across genera. In contrast, pairs with intermediate cointegration rates display higher variability.*

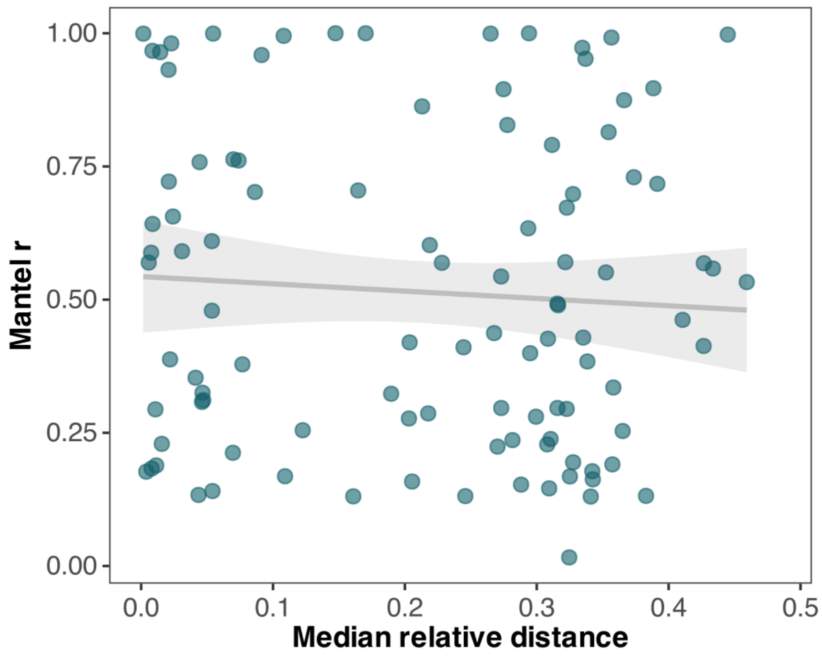


***Supplementary Figure 11. Relationship between median relative distance and phylogenetic congruence.*** *The scatter plot shows the association between the median relative distance and phylogenetic congruence (Mantel r) of each replicon pair. No significant association was detected (Spearman's $\rho = -0.05$, $p > 0.5$).*

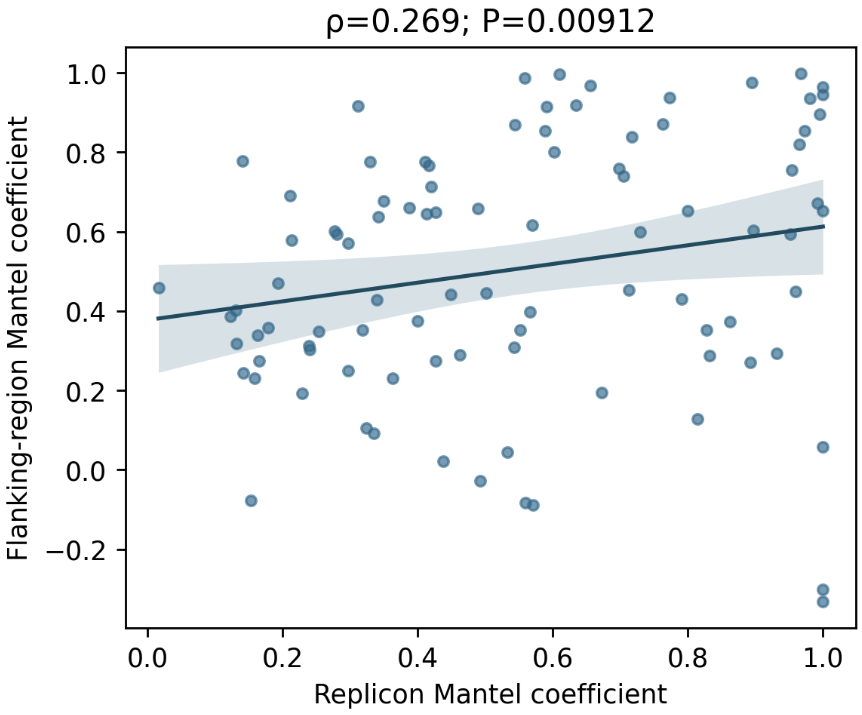


***Supplementary Figure 12. Association between replicon phylogenetic congruence and conservation of the surrounding genomic context.*** *Each point represents one replicon pair. The x-axis shows the Mantel correlation between replicon phylogenies, whereas the y-axis shows the Mantel correlation between their corresponding flanking regions. The grey line shows the linear fit. Spearman rho and p-value are shown above the plot.*

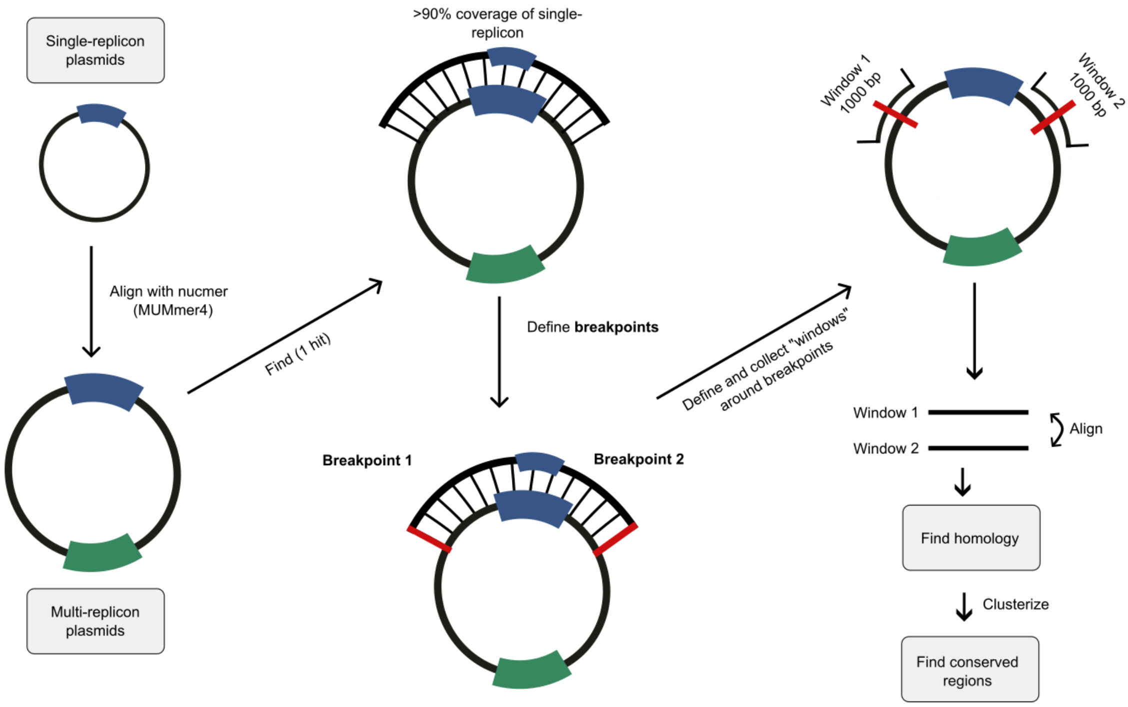


***Supplementary Figure 13. Workflow for the identification and characterization of plasmid fusion breakpoints and associated homologous regions.*** *Single-replicon plasmids are aligned against multireplicon plasmids using NUCmer from MUMmer4 to identify high-coverage matches (>90% coverage of the single replicon), enabling the detection of putative fusion events. For each event, the boundaries of the aligned region are used to define two fusion breakpoints on the multireplicon plasmid. Sequence windows centered on each breakpoint are then extracted and compared pairwise to identify homologous sequences. Detected homologous regions are subsequently clustered to define conserved sequence elements associated with fusion events.*

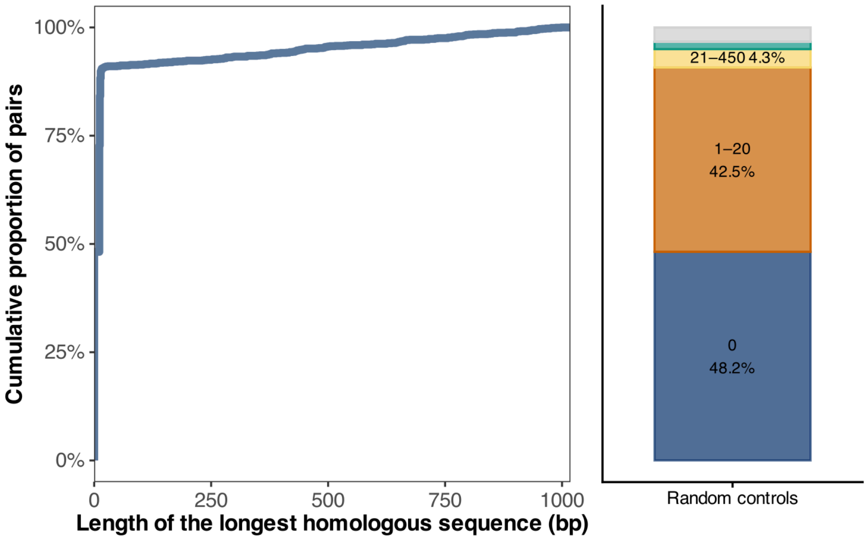


***Supplementary Figure 14. Distribution of homologous sequence lengths in random control breakpoint windows.*** *Cumulative distribution of the longest homologous sequence identified between pairs of random windows (±500 bp) sampled from the same plasmid set used in the fusion analysis (Figure 5). The stacked bar plot summarizes the proportion of window pairs by homology length categories. For each category, the upper value represents the base pair (bp) range, while the lower value shows the percentage of occurrences.*

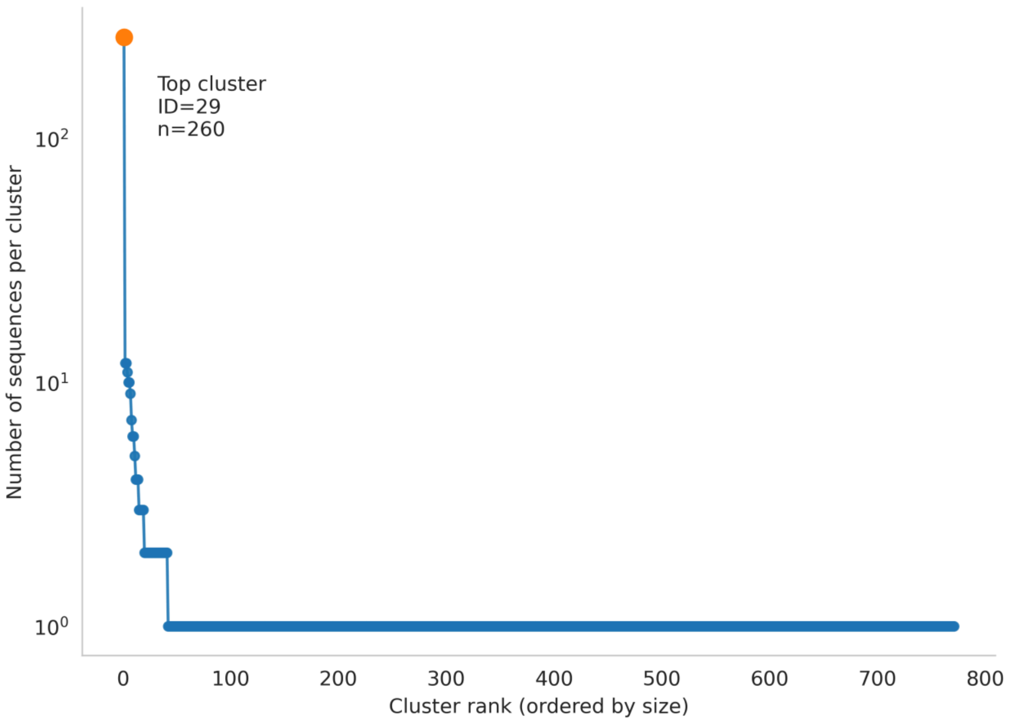


***Supplementary Figure 15. Rank–abundance distribution of homologous sequence clusters.*** *Clusters were ordered from largest to smallest according to the number of sequences they contain (cluster rank, x-axis). The y-axis shows cluster size on a log scale. The distribution is highly skewed, with a single dominant cluster and a long tail of low-prevalence clusters, indicating that most homologous tracts are rare while a small subset is widely shared.*

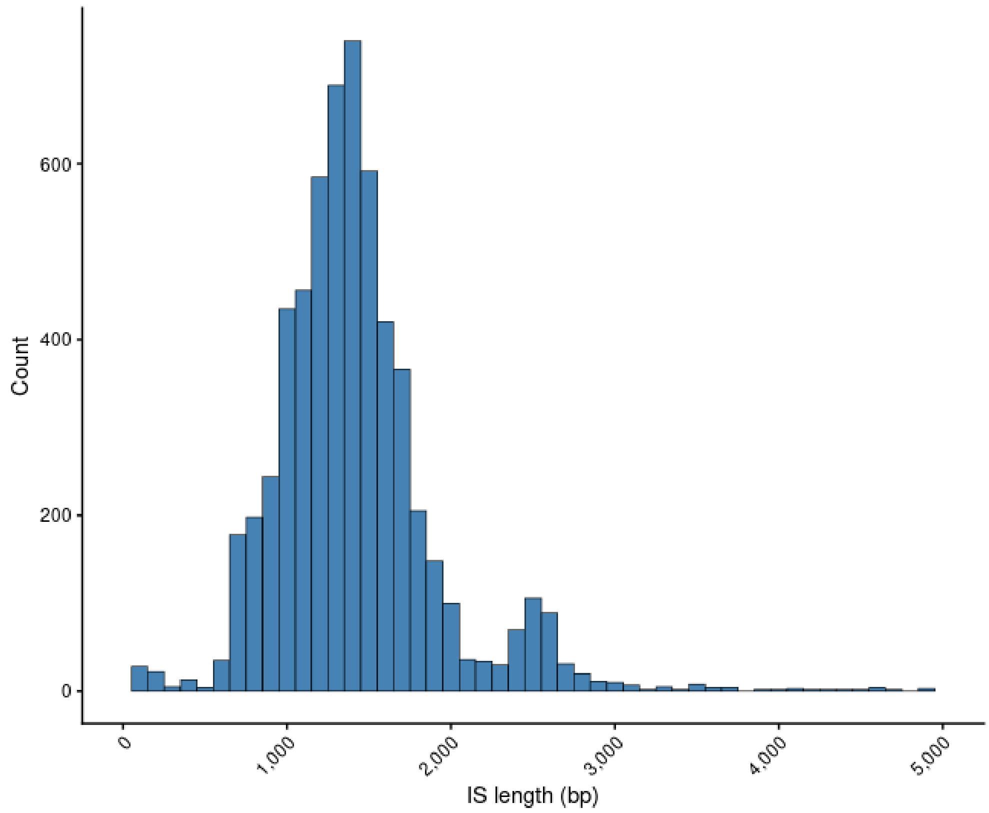


***Supplementary Figure 16. Length distribution of IS elements.*** *Bars show the raw counts (y-axis) of elements of given length (X-axis) in the ISfinder database.*

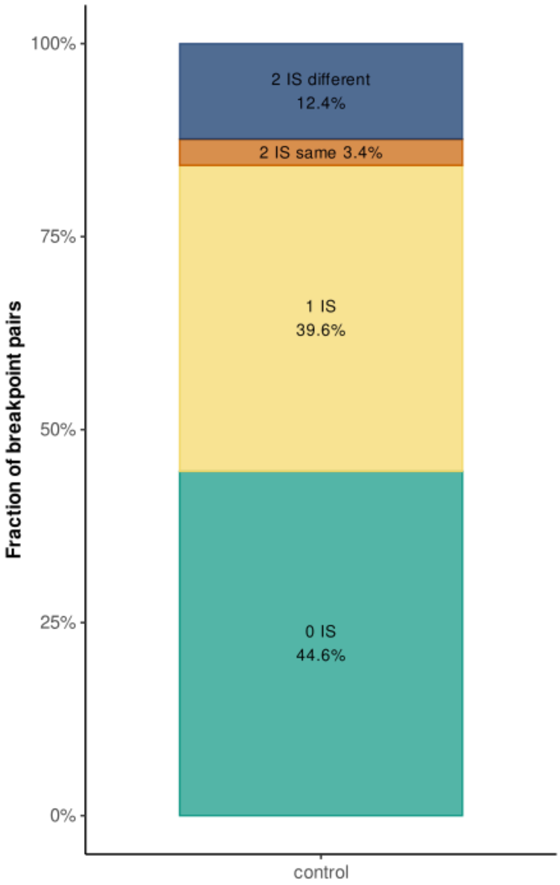


***Supplementary Figure 17. IS elements found in random windows.*** *The stacked bar shows the fraction of times 0 IS, 1 IS, 2 IS (duplicated), or 2 IS (different ones) are found when searching random windows of 4kb in the cointegrate plasmids that contain a cointegration event.*

**Supplementary Tables**

**Supplementary Table 1. Adjusted associations with multireplicon status for gene content**

| Category | Adjusted OR | 95% CI | P value |
|---|---|---|---|
| AMR genes | 1.78 | 1.65–1.90 | 1.98e-57 |
| Metal/biocide resistance genes | 1.77 | 1.62–1.92 | 1.54e-38 |
| Virulence genes | 4.40 | 3.73–5.20 | 5.33e-68 |

Separate binomial logistic regression models were fitted for AMR, metal/biocide resistance, and virulence gene presence, with multireplicon status as the focal predictor and adjustment for $\log_{10}$-transformed plasmid length, predicted mobility category, host genus, and isolation source.

**Supplementary Table 2. Adjusted associations with multireplicon status for mobility categories**

| Category | Adjusted OR | 95% CI | P value |
|---|---|---|---|
| pCONJ | 1.22 | 1.13–1.31 | 9.48e-08 |
| pMOB | 1.13 | 1.05–1.23 | 0.0015 |
| pOriT | 2.11 | 1.89–2.35 | 2.02e-41 |
| pNT | 0.50 | 0.46–0.53 | 4.93e-78 |

Separate binomial logistic regression models were fitted for each predicted mobility category (pCONJ, pMOB, pOriT, and pNT) versus all other categories, with multireplicon status as the focal predictor and adjustment for $\log_{10}$-transformed plasmid length, host genus, and isolation source.
